# Detection and typing of plasmids in *Acinetobacter baumannii* using *rep* genes encoding replication initiation proteins

**DOI:** 10.1101/2022.08.26.505409

**Authors:** Margaret M.C. Lam, Jonathan Koong, Kathryn E. Holt, Ruth M. Hall, Mehrad Hamidian

## Abstract

Plasmids found in *Acinetobacter* species contribute to the spread of antibiotic resistance genes. They appear to be largely confined to this genus and cannot be typed with available tools and databases. Here, a method for distinguishing and typing these plasmids was developed using a curated, non-redundant set of 621 complete sequences of plasmids from *Acinetobacter baumannii*. Plasmids were separated into three groups based on the Pfam domains of the encoded replication initiation (Rep) protein and a fourth group that lack an identifiable Rep protein. The *rep* genes of each Rep-encoding group (n=13 Rep_1, n=107 RepPriCT_1, n=351 Rep_3) were then clustered using a threshold of >95% nucleotide identity to define 80 distinct types. Five Rep_1 subgroups, designated R1_T1 to R1-T5, were identified and a sixth reported recently was added. Each R1 type corresponded to a conserved small plasmid sequence. The RepPriCT_1 plasmids fell into 5 subgroups, designated RP-T1 to RP-T5 and the Rep_3 plasmids comprised 69 distinct types (R3-T1 to R3-T69). Three R1, 2 RP and 32 R3 types are represented by only a single plasmid. Over half of the plasmids belong to the four most abundant types: the RP-T1 plasmids (n=97), which include conjugation genes and are often associated with various acquired antibiotic resistance genes, and R3-T1, R3-T2 and R3-T3 (n=95, 30 and 45, respectively). To facilitate typing and the identification of plasmids in draft genomes using this framework, we established the *Acinetobacter* Typing database containing representative nucleotide and protein sequences of the type markers (https://github.com/MehradHamidian/AcinetobacterPlasmidTyping).

**IMPORTANCE:** Though they contribute to the dissemination of genes that confer resistance to clinically important carbapenem and aminoglycoside antibiotics used to treat life-threatening *Acinetobacter baumannii* infections, plasmids found in *Acinetobacter* species have not been well studied. As these plasmids do not resemble those found in other Gram-negative pathogens, available typing systems are unsuitable. The plasmid typing system developed for *A. baumannii* plasmids with an identifiable *rep* gene will facilitate the classification and tracking of sequenced plasmids. It will also enable the detection of plasmid-derived contigs present in draft genomes that are widely ignored currently. Hence, it will assist in the tracking of resistance genes and other genes that affect survival in the environment, as they spread through the population. As identical or similar plasmids have been found in other *Acinetobacter* species, the typing system will also be broadly applicable in identifying plasmids in other members of the genus.

## INTRODUCTION

The plasmids found in *Acinetobacter* species clearly differ from the better studied and understood plasmids found in the majority of Gram-negative species and covered by the PlasmidFinder database (1). Indeed, the plasmids found in other Gram-negative species (especially Enterobacterales) do not appear to be stably maintained in *Acinetobacter* species and *Acinetobacter* plasmids are not seen in other Gram-negative pathogens. Hence, a typing and classification scheme relevant to *Acinetobacter* plasmids is needed.

In 2010, sequences were available for very few *Acinetobacter* plasmids, and they were mainly derived from the modest number of complete genomes of *A. baumannii* isolates that had been published at that time. At this point, an analysis aimed at generating a PCR detection and typing scheme based on *rep* genes encoding replication initiation (Rep) proteins was published (2). Fifteen complete plasmid sequences, 12 of them derived from 5 available complete genomes, together with 8 partial sequences (5 determined in the study) were analysed. Four plasmids were identical or had identical or very closely related *rep* genes, and two plasmids included two and another has three *rep* genes. Hence, a total of 24 different *rep* gene sequences were examined. The majority encoded Rep proteins belonging to Pfam01051 corresponding to the Rep_3 family. The Rep proteins of two plasmids belonged to the Rep_1 group (Pfam01446) and one plasmid encoded a protein with a Rep motif (Pfam03090). As the method developed used PCR to detect the *rep* genes, a cut off of 74% nucleotide identity in the *rep* gene sequence was used to group the *rep* genes and ensure specificity of the primers. Nineteen groups (GR1-GR19) were proposed. However, some of these groups encompassed two clearly distinct *rep* gene types and a secondary classification assigned Aci numbers up to 10 to some of the Rep proteins, e.g. the replicases associated with the two distinct groups in GR2 were designated Aci1 and Aci2 (2).

Using the PCR approach, an analysis of 96 multiply antibiotic resistant isolates, mainly from Europe, by the same group found that *rep*Aci1 (GR2) and *rep*Aci6 (GR6) were the most widely distributed (3). However, as the cost of genome sequencing fell, over the next few years many more complete or draft genomes became available and PCR typing was never extensively used. Consequently, such broad groups including members with *rep* genes that differ by up to 26% nucleotide identity are no longer necessary or practical. Moreover, in the absence of either a centralised database or a clear revision of the rules for grouping, assignment of additional groups has been problematic. For example, the GR20 designation has been used for at least three different types (4–6). In other cases, examples of new types have been identified but GR numbers were not assigned (e.g (7, 8).

In 2017, a review examined only the small (<10 kbp) *Acinetobacter* plasmids and, based on a phylogeny of the Rep_3 group proteins, separated plasmids encoding the distinct Aci1 and Aci2 types of the GR2 group with the Aci2 type becoming GR20 (5). Though this separation breaches the 74% rule as these *rep* genes are close to 80% identical, it appears to have prevailed in more recent studies (see below). However, the distinct subtypes found in other GR were not separated and additional groups were not identified. The number of groups was expanded to 23 in a 2017 study that compared three plasmids from a single *A. baumannii* isolate from Argentina to a database of 122 Rep proteins found in *A. baumannii* plasmids (9). A cut-off of 85% protein identity was used to split GR8 into two groups, GR8 and GR23, and two further GR (GR21 and GR22) were reported (9). In 2020, a comprehensive analysis of complete plasmids from *A. baumannii* available in 2016 added several further GR up to GR33 using a cut off of >74% nucleotide identity (10). Finally, in 2021, GR34 was assigned, again based on the original cut off of >74% DNA identity (11).

A shortcoming of the GR scheme for *rep* genes arises from the fact that a number of plasmid types found in *A. baumannii* and in other *Acinetobacter* species do not include an identifiable *rep* gene and some of these are common (12–14). To overcome this, a few further studies have attempted to address the issue of *Acinetobacter* plasmid classification using various different data sets (new sequences and published sequences) and different approaches. Salto and co-workers (15) looked at replication, mobilisation and conjugative transfer genes in a group of plasmids from various *Acinetobacter* species. They also identified 16 groups of Rep_3 family proteins that were designated AR3G1–AR3G16 from plasmids recovered from various species but the numbering of these AR3 groups does not correspond to the group numbers used by Bertini *et al*. and a key was not provided. In 2020, Mindlin and co-workers (14) published a study of plasmids isolated from *Acinetobacter lwoffii* isolates recovered from permafrost, where they used a classification system for plasmids from any *Acinetobacter* species based on size, then whether a *rep* or a *mob* gene, both or neither was present. They used the Salto *et al*. groupings for the Rep_3 proteins and a cut-off of 95% protein identity for clustering.

The study of Salgardo Camargo and co-workers (10) examined 18 new sequences and 145 *A. baumannii* plasmids for which complete sequences were available in 2016 and grouped them using an approach that attempted to classify the whole plasmids into lineages without accounting for the many known accessory regions that should be removed to reveal the plasmid backbone. Hence, their approach is confounded by the extent of the variation caused by the acquisition and loss of significant portions of plasmids with closely-related backbones, as occurs for example in the small to medium sized Rep_3 plasmids that carry acquired *dif* modules where the size of the *dif* module and other accessory content can exceed that of the backbone (e.g. (7, 16–18). They retained the >74% nucleotide identity in *rep* genes cut off used by Bertini and co-workers (2) and removed GR23 but kept GR20. They added 10 GR bringing the total to 33 GR. Nine of the additional plasmid types encoded Rep, Rep_1 and Rep_3 replication initiation proteins; the tenth, represented by a single plasmid, encoded a protein with a RepC motif that is not involved in replication of the plasmid (see below). A phylogeny of the Rep_3 proteins revealed distinct subgroups within some GR, as was found previously (2).

Here, we have analysed the plasmids from *A. baumannii* with a complete sequence available in the GenBank nucleotide database as of February 2021 in order to develop a unified system that will allow plasmids to be typed simply and rapidly and to facilitate identification of plasmids in draft genome sequences. To reduce the plasmid data set to a manageable size, only *A. baumannii* plasmids were included as these are the most important from a clinical perspective and, as a first step, only those encoding a Rep protein have been typed. The most appropriate criteria for defining plasmid groups based on *rep* genes was re-assessed in the light of the fact that PCR is no longer the main source of information about the types of plasmids carried by *A. baumannii* isolates. A new typing and numbering system was developed but for ease of comparison, the earlier GR designations are indicated, where relevant. An online resource that includes a database of representative *rep* gene and Rep protein sequences was developed and has been made available via GitHub.

## RESULTS

### Curation of the plasmid database

All complete sequences for *A. baumannii* associated plasmids were downloaded (February 2021) and, after removal of redundancies, further curation removed additional plasmids from the data set (see Methods and reasons listed in Supplementary Table S1). Of particular note, sequences that were previously classified as lineage 10 (10) and assigned to GR3 were removed as they have been shown to represent a circular form of the AbGRI3 resistance island that has failed to assemble to the correct location on the chromosome (19). The segment includes only an incomplete and inactive *rep* gene (see (20) for details). The final data set comprised 621 plasmids, 539 of them derived from genome sequencing projects. A small number of partial sequences reported by Bertini *et al*. (2) were also included in the analysis to facilitate correlation with the typing scheme devised by those authors but were not included in plasmid counts.

### Detection of *rep* genes

A comprehensive approach to detection of *rep* genes found in the plasmids in the data set (see Methods) included curation to remove genes incorrectly annotated as *rep* genes (described in more detail below). A total of 142 plasmids (23%) did not encode an identifiable Rep protein. Many of the plasmids in this group were related to known plasmids, including n=27, 7 and 20 identical to the well characterised, small plasmids pRAY* (13), pD36-1 (17), pA85-1b (21). Thirty one had backbones related to the backbones of large conjugative plasmids where a *rep* gene of a novel, as yet unidentified, type may be present. Among these, n=20 were related to pAB3 and pA297-3 (12), 8 related to pNDMBJ01 (22) and 3 related to pALWED1.1 (23). Investigation of the RepC protein encoded only by the plasmid pAB3 that was used to define GR33 (10), revealed that it is not found in known relatives with closely-related backbones such as pA297-3 (12) and it was traced to the genomic island *GIsul2* (24) which is found in pAB3 but not in its close relatives. As the related plasmids do not include an identifiable *rep* gene, pAB3 was included in the no *rep* plasmid group. This highlights the importance of careful curation to identify problems that arise when bioinformatic approaches are used without reference to underlying knowledge of which regions are plasmid backbone and which are accessory. The remaining (n=46) plasmids with no identifiable *rep* gene were not further examined in this study.

At least one *rep* gene was identified in 479 plasmids (77%) (Table 1). However, 19 plasmids included 2 *rep* genes and 2 included 3 *rep* (Table 1), yielding in total 502 *rep* genes. The product of each *rep* gene was screened for Pfam domains associated with replication initiation (Rep) proteins (see Methods).

**TABLE 1.** Summary of plasmid sequence data studied

| Plasmids | Number |
| --- | --- |
| Complete plasmids | **621** |
| Part of Genomes | 539 |
| Not part of a genome project | 82 |
| No Rep | 142 |
| With Rep<sup>a</sup> | 479 |
| Plasmids encoding Rep_3 | 359 |
| Plasmids encoding Rep_1 | 13 |
| Plasmids encoding RepPriCT_1 | 107 |
| Total plasmids | **479** |

<sup>a</sup> 457, 19 and 2 plasmids encode 1, 2, and 3 Reps, respectively.

### Classification of plasmids carrying *rep* genes

To simplify classification we have used the Pfam group of the replication initiation protein encoded by each *rep* gene for initial grouping of the plasmids, as this can be obtained readily. Plasmids encoding a Rep_1 family (Pfam01446) replicase were least abundant with only 13 plasmids in this group (Table 1). The Rep_3 (Pfam01051) group predominated with 359 plasmids and all of the plasmids carrying multiple *rep* genes fell into this category. The Rep group (Pfam03090) that encode replicases that also include a PriCT_1 (Pfam08708) motif, here referred to as the RepPriCT_1 group, included 107 plasmids. The distinct types within each Pfam group were then defined by clustering the *rep* nucleotide sequences (see Methods), using a cut-off of 95% nucleotide identity as this appeared to best separate the clearly distinct types without recording minor variations in DNA sequence. However, for most types identified using this approach, all represented *rep* genes were >99% identical.

A total of 80 types were identified using this approach: 6 Rep_1 types, 69 Rep_3 types and 5 RepPriCT_1 types. The plasmid types identified in this way have been prefaced R1, R3 and RP for the Rep_1, Rep_3 and RepPriCT_1 groups, respectively, followed by an assigned number, generally in the order of identification or the relative abundance of the type. To facilitate comparison to earlier studies, where relevant the GR number is indicated in Tables.

### The small Rep_1 plasmids

In the original plasmid classification (2), one plasmid, p4ABAYE (GR14) and a partial sequence from pAB49 (GR16) encoded a Rep protein belonging to the Rep_1 family (Pfam01446). The complete sequence of pAB49 is now available (GenBank accession number L77992.1) and, though it was not detected in our initial search for complete plasmids, was added to the complete plasmid database.

Only 11 additional Rep_1 plasmids were found among the 621 plasmids examined here. Of these, the complete sequences of seven were identical to either p4ABAYE or pAB49 and these groups were designated R1-T1 and R1-T2, respectively (Table 2). Within each type, very little variation in the sequences was observed indicating that these plasmids are well conserved. These types were also widely distributed as they were recovered in various countries (Table 2). Three additional types (R1-T3 to R1-T5), each represented by only one or two examples, were detected. An additional Rep_1 plasmid type was reported recently (25) and has been added to Table 2 as R1-T6.

**TABLE 2.** Properties of plasmids encoding Rep_1 (Pfam01446) family replication initiation protein.

| **Rep type** | **Plasmid name** | **Length (bp)** | **Strain name** | **Year** | **Country** | **GenBank Acc. no.** | **Rep Protein id** |
| --- | --- | --- | --- | --- | --- | --- | --- |
| R1-T1 | **p4ABAYE<sup>a</sup>** | 2726 | AYE | 2001 | France | CU459139.1 | CAM84626.1 |
| R1-T1 | pA85-1 | 2726 | A85 | 2003 | Australia | CP021783.1 | AHM95258.1 |
| R1-T1 | pMRSN58-2.7 | 2725 | MRSN 58 | 2010 | USA | CM003316.1 | KLT88566.1 |
| R1-T1 | p4ZQ6 | 2725 | ZQ6 | 2016 | Iraq | CM009041.2 | PQL72120.1 |
| R1-T1 | p4ZQ5 | 2725 | ZQ5 | 2016 | Iraq | CM009683.1 | PQL83905.1 |
| R1-T2 | **pAB49<sup>b</sup>** | 2343 | Ab49 | nk | nk | L77992.1 | AAA99423.1 |
| R1-T2 | pA85-1a | 2343 | A85 | 2003 | Australia | CP021784.1 | ASF79212.1 |
| R1-T2 | pMRSN7339-2.3 | 2343 | MRSN 7339 | 2004 | USA | CM003313.1 | KLT74217.1 |
| R1-T2 | pDA33382-2 | 2359<sup>d</sup> | DA33382 | nk<sup>a</sup> | Germany | CP030108.1 | AXB17575.1 |
| R1-T3 | pAb-D10a-a_5 | 2697 | Ab-D10a-a | 2016 | Ghana | CP051874.1 | QJF33731.1 |
| R1-T3 | pAb-B004d-c_4 | 2697 | Ab-B004d-c | 2016 | Ghana | CP051879.1 | QJF37612.1 |
| R1-T4 | p3AB5075 | 1967 | AB5075-UW | 2008 | USA | CP008709.1 | AKA33693.1 |
| R1-T5 | pTS236 | 2252 | DS002 | nk | nk | JN872565.1 | AFB69861.1 |
| R1-T6 | pMRSN56-1 | 2178 | MRSN56 | 2010 | USA | CP080453.1 | QYM41639.1 |

<sup>a</sup> GR14 in (2).<sup>b</sup> GR16 sequence was reported as partial in (2) but completed later in 2016.<sup>c</sup> not known.<sup>d</sup> length difference is likely due to an assembly issue.

Plasmids with a Rep_1 replication initiation protein are generally small and replicate using a rolling circle mechanism. The Rep_1 plasmids identified here were all less than 3 kbp in length and do not include any antibiotic resistance genes. Most were found in completed genome sequences. However, due to their small size, they may be missed in genomes derived using long read sequencing (26). The relationship between the sequences of the Rep proteins encoded by each type is shown in Table 3. There appear to be two distinct groups based on alignments with significant levels of identity and >75% coverage, namely T1 and T5 in one group and T2, T3, T4 and T6 in the other. The structures of one plasmid of each type in the R1 group are shown in Figure 1.

**FIG 1.**
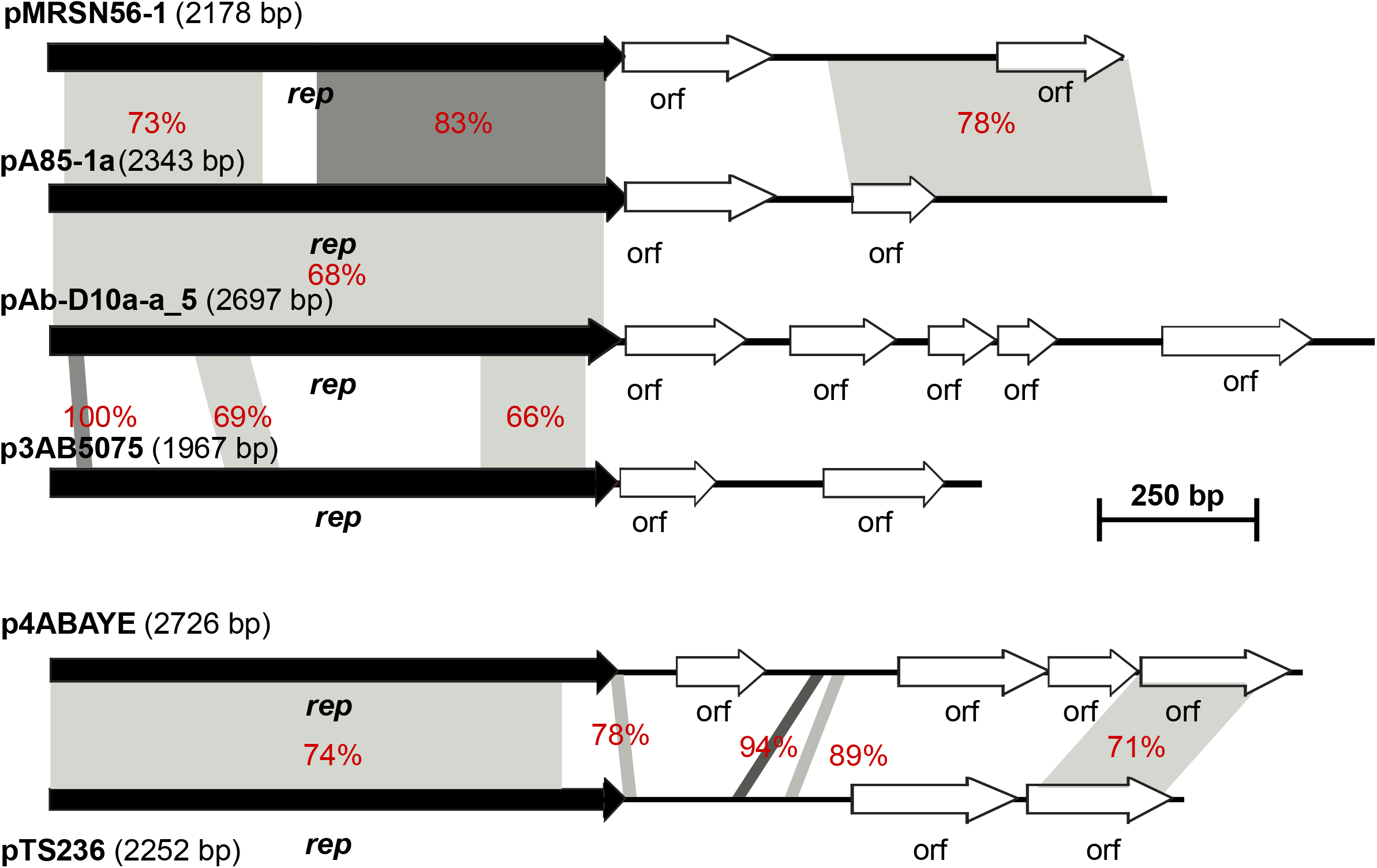
Comparison of representatives of plasmids encoding Rep_1 family (Pfam01446) Rep proteins. Black horizontal arrows show indicate *rep* genes. White arrows encode hypothetical proteins. Regions with significant DNA identities are shown using shades of grey with % identities also labelled in red. p4ABAYE (GenBank accession number CU459139), pA85-1a (GenBank accession number CP021784), pAb-D10a-a_5 (GenBank accession number CP051874), p3AB5075 (GenBank accession number CP008709), pTS236 (GenBank accession number JN872565), and pMRSN56-1 (GenBank accession number CP080453) represent R1-T1, R1-T2, R1-T3, R1-T4, R1-T5 and R1-T6, respectively.

**TABLE 3.** Pairwise protein identities of representative Rep_1 type Rep proteins ^a^

| Type | P1-T1 | P1-T2 | P1-T3 | P1-T4 | P1-T5 | P1-T6 |
| --- | --- | --- | --- | --- | --- | --- |
| P1-T1 | 100 | 25.2 (27) | 28.2 (31) | - <sup>b</sup> | 73.4 (82) | 26.68 (36) |
| P1-T2 |  | 100 | 63.7 (89) | 44.4 (82) | 22.7 (28) | 73.28 (100) |
| P1-T3 |  |  | 100 | 45.2 (76) | 26.7 (30) | 65.63 (90) |
| P1-T4 |  |  |  | 100 | - | 44.58 (82) |
| P1-T5 |  |  |  |  | 100 | 24.8 (28) |
| P1-T6 |  |  |  |  |  | 100 |

<sup>a</sup> numbers in bracket indicate % coverage.
<sup>b</sup> no significant matches.

### The RepPriCT_1 family plasmids

In the original classification, a single plasmid pACICU2, carried a gene encoding a Rep protein belonging to the Rep and PriCT_1 families. The plasmid type was GR6 and the Rep protein was designated Aci6 (2). Plasmids of this type have been implicated in the dissemination of the *oxa23* gene (carbapenem resistance) and the *aphA6* gene (amikacin resistance). They include a complete set of genes for conjugation, and some have been shown to be conjugative (27–29). Among the 107 complete RepPriCT_1 plasmids analysed here, the majority (n=97) were *rep*Aci6 plasmids, here designated RP-T1. However only 10 representatives are listed in Table 4 selected to include well characterised examples and to illustrate the global distribution of plasmids of this type. A complete list can be found in Supplementary Table S2. Where it has been examined, these plasmids share a common, though not completely identical, backbone that is often interrupted by transposons encoding either antibiotic resistance genes or heavy metal resistance genes (28–31). Hence, they are one of the most important plasmid types implicated in introducing further antibiotic resistance genes into *A. baumannii* isolates.

**TABLE 4.** Properties of plasmids encoding Rep family (Pfam03090) replication initiation protein

| Rep type<sup>a</sup> | Plasmid | length (bp) | Date | Country | Strain name | Accession number | Rep Protein id |
| --- | --- | --- | --- | --- | --- | --- | --- |
|  | name |  |  |  |  |  |  |
| RP-T1 | pACICU2<sup>b</sup> | 70101 | 2005 | Italy | ACICU | CP031382 | QCS04021 |
| RP-T1 | pAb-G7-2<sup>b</sup> | 70100 | 2002 | Australia | G7 | KF669606 | AGY56182 |
| RP-T1 | pA85-3<sup>b</sup> | 86,334 | 2003 | Australia | A85 | CP021787 | ASF79226 |
| RP-T1 | pS21-2<sup>b</sup> | 123432 | 1996 | Singapore | SGH9601 | MG954377 | AWO68424 |
| RP-T1 | pMC23.1 | 67441 | 2016 | Bolivia | MC23 | MK531538 | QCO89676 |
| RP-T1 | pCS01A<sup>c</sup> | 63720 | <2014 | Switzerland | CS01 | HG977523 | -<sup>c</sup> |
| RP-T1 | pTG22182_1 | 71233 | 2011 | USA | TG22182 | CP039994 | QCO84474 |
| RP-T1 | p3VB958 | 82500 | 2019 | India | VB958 | CP040043 | QCP18569 |
| RP-T1 | pAB04-2 | 87569 | 2012 | Canada | Ab04-mff | CP012008.1 | AKQ32652.1 |
| RP-T1 | p2K50 | 79598 | 2008 | Kuwait | K50 | LT984690 | SPC58401 |
| RP-T2 | pABTJ1<sup>d</sup> | 77528 | nk<sup>e</sup> | China | MDR-TJ | CP003501 | AFI97405 |
| RP-T2 | pAZJ221<sup>f</sup> | 77530 | 2009 | China | A221 | KM922672 | AJF79875 |
| RP-T2 | p1BJAB07104 | 70170 | 2007-8 | China | BJAB07104 | CP003887 | AGQ16162 |
| RP-T2 | p2BJAB0868 | 70167 | 2007-8 | China | BJAB0868 | CP003888 | AGQ12301 |
| RP-T2 | pHRAB-85 | 77513 | 2014 | China | HRAB-85 | CP018144 | APF45706 |
| RP-T2 | pBJ83 | 69069 | 2007 | China | XDR-BJ83 | CP018422 | APM50974 |
| RP-T2 | pABAY10001_1C | 54627 | <2019 | South Korea | ABAY10001 | MK386682 | QBN23313 |
| RP-T3 | pKBN10P02143<sup>g</sup> | 52517 | 2012 | South Korea | KBN10P02143 | CP013925 | ALY01450 |
| RP-T4 | p2ZQ6 | 6772 | 2016 | Iraq | ZQ6 | CM009039 | PQL72098 |
| RP-T5 | p4.36-1512 | 4721 | 2016 | Russia | 36-1512 | CP059390 | QLY88352 |
| RP-T5 | pE47_007 | 4715 | 2013 | Australia | E47 | CP042563 | QFH47718 |

<sup>a</sup> 94 total plasmids including pAbG7-2, pA85-3, pD46-3, pS32-2 have a Rep protein greater than 99% identical to Rep encoded by pACICU2. Ten representatives are shown, and a complete list is in Supplementary Table S2. This type corresponds to GR6.
<sup>b</sup> shown to conjugate (16, 27-29).
<sup>c</sup> A second plasmid pCS01B is identical to pCS01A but the *rep* gene is incomplete. Entry does not include annotations and hence no protein id.
<sup>d</sup> pABTJ1 is grouped as GR25 in (10).
<sup>e</sup> not known.
<sup>f</sup> shown to be conjugative (6). As for pABTJ1 it carries Tn2009 (containing *oxa23*).
<sup>g</sup> pKBN10P02143 is designated as GR32 (10).

A group of 7 plasmids were close relatives of pABTJ1 (32) and this type, here designated RP-T2, corresponds to GR25. The backbone of plasmid pABTJ1 has been shown to include a set of genes that encode proteins involved in conjugation that are related to those encoded by the RP-T1 (Aci6) group (33). Another plasmid of this type also carrying the carbapenem resistance transposon *Tn2009* has been shown to be conjugative (6). These plasmids are so far confined to Asia (Table 4). Three further types, RP-T3 to RP-T5 were each detected in only one or two plasmids (Table 4) with RP-T3 corresponding to GR32, and no reports describing them were found.

The relationships between the Rep proteins encoded by the types in this group are shown in Table 5. This comparison revealed two broad subgroups, with RP-T1 Rep grouping with RP-T2, and RP-T4 grouping with RP-T5 while RP-T3 was more distantly related to the other types. RP-T4 and RP-T5 plasmids are substantially smaller than the conjugative RP-T1 and RP-T2 plasmids.

**TABLE 5.** Pairwise protein identities of Rep group representatives ^a^

| Rep type | Plasmid | Protein id | pACICU2 | pABTJ1 | pKBN10P02143 | p2ZQ6 | p4.36-1512 |
| --- | --- | --- | --- | --- | --- | --- | --- |
| RP-T1 | pACICU2 | QCS04021.1 | 100 | 69.23 (98) | 38.7 (95) | 38.6 (83) | 37.1 (85) |
| RP-T2 | pABTJ1 | AFI97405.1 |  | 100 | 39.7 (89) | 40.8 (79) | 36.6 (80) |
| RP-T3 | pKBN10P02143 | ALY01450.1 |  |  | 100 | 35.9 (98) | 35.2 (71) |
| RP-T4 | p2ZQ6 | PQL72098.1 |  |  |  | 100 | 69.3 (100) |
| RP-T5 | p4.36-1512 | QLY88352.1 |  |  |  |  | 100 |

<sup>a</sup> numbers in bracket indicate % coverage.

### Identification of the *rep* gene in Rep_3 family plasmids

There has significant confusion in the literature and in the annotations of plasmid sequences in GenBank with respect to the identification and naming of the *rep* gene in many of the plasmids in this category. The confusion applies particularly to plasmids with the configuration shown in Figure 2, and complicated the identification of *rep* genes for inclusion in the *rep* gene database. The confusion appears to have arisen early when the *rep* gene was designated *repB* and the adjacent downstream gene, which we have labelled orfX in Figure 2 as its function is currently unknown, was called *repA* (34). In fact, a published study had previously demonstrated that only a single gene (the one designated *repB*) and upstream iterons are essential for replication of pMAC (p2ATCC19606) and the *rep* gene, which encodes the Aci9 replicase (2), was designated *repM* (35). The presence of orfX, which encodes a helix-turn-helix domain protein, in the plasmid set used by Bertini and co-workers is also shown in Table 6. Currently, the downstream orfX gene continues to be misidentified in many publications and GenBank entries as the *rep* gene, and manual curation (see Methods) was used to ensure that these genes were not included in our databases.

**FIG 2.**
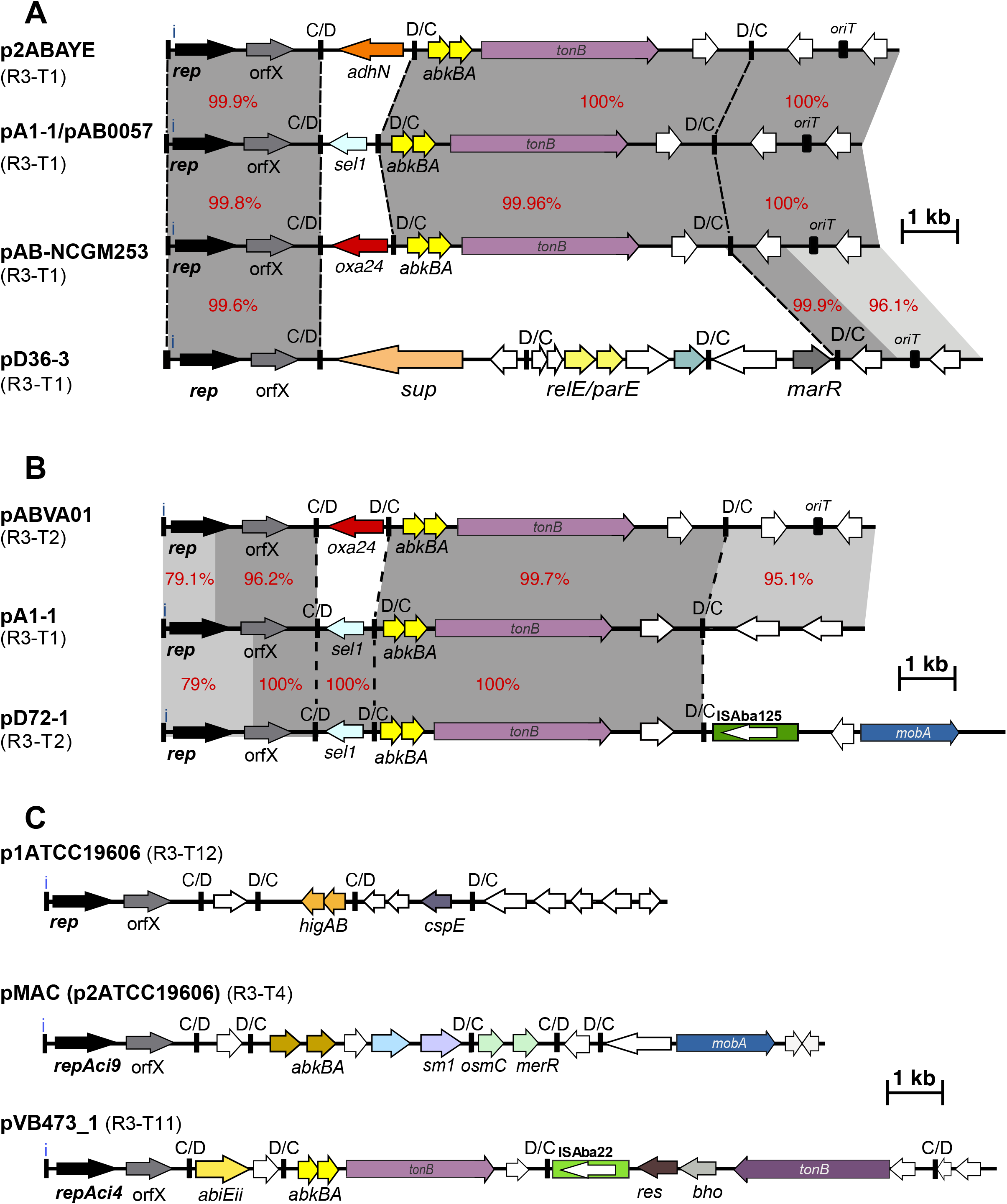
Schematic comparison of Rep_3 family (Pfam01051) plasmid structures. Horizontal arrows show the length and orientation of genes with *rep* genes coloured black, resistance genes red, toxin/anti-toxins yellow and mobilisation genes blue. Green boxes indicate insertion sequences with their transposase shown inside the box. Small thick vertical bar marked with “i” indicate iterons. Other vertical bards marked with “C/D or D/C” indicate the location of p*dif* sites. Regions with significant DNA identities are shown using shades of grey with % identities also shown using red numbers. Dotted lines draw the show the boundaries of p*dif* modules. Panel A represents variations within five R3-T1 plasmidS (p2ABAYE, pA1-1/pAB0055, pAB-NCGM253 and pD36-3 with GenBank accession numbers CU459138, CP010782/CP001183, AB823544, and CP012955, respectively) that carry different p*dif* modules. Panel B compares two R3-T1 plasmids pABVA01 and pD72-1 (GenBank accession numbers FM210331.1 and KM051986, respectively) with pA1-1 (GenBank accession number CP010782) representing R3-T1. Panel 3 represents the structure of plasmids representing R3-T12 (p1ATCC19606; GenBank accession number CP045108), R3-T4 (pMAC; AY541809) and R3-T11 (pVB473_1; GenBank accession number CP050389) with no significant DNA identity.

**TABLE 6.** Properties of plasmids encoding Rep_3 family proteins described in Bertini *et al*, 2010

| Plasmid | Rep | GR Group | Length (aa) | Iterons | OrfX | C/D | Protein_id | Plasmid length (bp) | Strain | accession no. | number complete 95% (99%) |
| --- | --- | --- | --- | --- | --- | --- | --- | --- | --- | --- | --- |
| pAB0057 | Aci1 | 2 | 316 | 4 | Y | Y | ACJ43223.1 | 8731 | AB0057 | CP001183 | 95 (88) |
| p2ABAYE | Aci1 | 2 | 316 | 4 | Y | Y | CAM84615.1 | 9661 | AYE | CU459138 | “ |
| pACICU1 | Aci1 | 2 | 316 | 4 | Y | Y | ACC58980.1 | 28279 | ACICU | CP000864 | “ |
| pAB2 | Aci1 | 2 | 316 | 4 | Y | Y | ACC58980.1 | 11302 | ATCC17978 | CP000523 | “ |
| pABVA01 | Aci2 | 2 | 316 | 4 | Y | Y | CAR65318.1 | 8963 | VA-566/00 | FM210331 | 30 (29) |
| p203 | Aci3 | 3 | 307 | 4 | Y<sup>a</sup> | Y<sup>a</sup> | ADM89092.1 | > 1068 | Ab203 | GU978997 | 13 (13) |
| p844 | Aci4 | 4 | 307 | 4 | Y<sup>b</sup> | Y<sup>b</sup> | ADM89093.1 | >1119 | Ab844 | GU978998 | 8 (8) |
| p537 | Aci5 | 5 | 310 | 4 | Y<sup>c</sup> | Y<sup>c</sup> | ADM89094.1 | >1125 | Ab537 | GU978999 | 2 (0) |
| pAb736 | Aci7 | 3 | 306 | 4 | Y<sup>d</sup> | Y<sup>d</sup> | ADM89091.1 | >1065 | Ab736 | GU978996 | 1 (0) |
| p11921 | Aci8 | 8 | 307 | 4 | N<sup>e</sup> | N<sup>e</sup> | ADM89095.1 | >1103 | Ab11921 | GU979000 | 2 (2) |
| pMAC | Aci9 | 8 | 311 | 4 | Y | Y | AAT09649.1 | 9540 | ATCC19606 | AY541809 | 15 (13) |
| pACICU1 | Aci10 (AciX) | 10 | 315 | 4 | N | N | ACC58984.1 | 28279 | ACICU | CP000864 | 4 (4) |
| p1ABAYE | p1ABAYE0001 | 11 | 316 | 4 | Y | Y | CAM84608.1 | 5644 | AYE | CU459137 | 4 (4) |
| pABIR | RepA_AB | 12 | 295 | 4<sup>f</sup> | Y | Y | ACB05788.1 | 29823 | Ab1 | EU294228 | 10 (1) |
| pAB02 | Rep_AB | 12 | 295 | 4 | Y | Y | AAR00517.1 | >4162 | Ab02 | AY228470 | 10 (9) |
| p3ABAYE | p3ABAYE0002 | 13 | 390 | - | N | N | CAM84695.1 | 94413 | AYE | CU459140 | 10 (7) |
| p1ABSDF | p1ABSDF0001 | 1 | 314 | 4 | N | N | CAP02936.1 | 6106 | SDF | CU468231 | 1 (1) |
| p2ABSDF | p2ABSDF0001 | 12 | 297 | 4<sup>f</sup> | Y | Y | CAP02944.1 | 25014 | SDF | CU468232 | 8 (8) |
| p2ABSDF | p2ABSDF0025 | 18 | 339 | 4 | N | N | CAP02966.1 | 25014 | SDF | CU468232 | 1 (1) |
| p3ABSDF | p3ABSDF0018 | 15 | 303 | 4<sup>f</sup> | Y | Y | CAP02992.1 | 24922 | SDF | CU468233 | 1 (1) |
| p3ABSDF | p3ABSDF0002 | 7 | 310 | - | N | N | CAP02976.1 | 24922 | SDF | CU468233 | 1 (1) |
| p3ABSDF | p3ABSDF0009 | 9 | 326 | 4<sup>f</sup> | N | N | CAP02983.1 | 24922 | SDF | CU468233 | 1 (1) |
| pAB1 | A1S_3471 | 17 | 318 | 4 | Y | Y | ABO13860.1 | 13408 | ATCC17978 | CP000522 | 1 (1) |
| p135040 | 135040 | 19 | 318 | 4 | nk | nk | ACX70400.1 | >3975 | 135040 | GQ861437 | 0 (0) |

<sup>a</sup> p203 is incomplete however another plasmid with an identical RepAci3 (pD1279779; GenBank no. CP003968) includes orfX and C/D sites downstream of the Rep.
<sup>b</sup> p844 is incomplete, (the RepAci4 plasmid pPM194122\_2; GenBank no. CP050427) includes orfX and C/D sites downstream of the Rep.
<sup>c</sup> p537 is incomplete but plasmid II of the strain R2091 (GenBank no. LN997847) with 98.6% DNA identity (compared to *repAci5* of p537) was analysed.
<sup>d</sup> pAb736 is incomplete however, the closet plasmid is pRCH52-1 (GenBank no. KT346360), *repAci7* and is 95% DNA identical to that of pRCH52-1.
<sup>e</sup> Given that p11921 is incomplete the presence of orfX and its downstream C/D sites were investigated in pAb825\_36 (GenBank no. MG100202), which encodes
a Rep identical to p11921. pAb825\_36 has iteron sequences upstream of its two *rep* genes.
<sup>f</sup> Imperfect iterons (1-3 bp differences).

### Overview of the Rep_3 family

Rep_3 plasmids were the most abundant in this collection and constitute the most diverse group. Classification of the 382 *rep* genes encoding Rep_3 replication initiation proteins using the cut-off of >95% nucleotide identity for the *rep* gene revealed 69 distinct types (Table 7). A complete list of the members of each type is found in Supplementary Table S3. However, only 8 types (T1-T8) included 10 or more members (accounting for 63% of *rep* genes encoding a Rep_3 type protein) and the majority of types are represented by only 1-4 members. When only representatives in the Bertini *rep* gene set are considered but a 95% nucleotide identity cut off is used more than half of the *rep* genes in the larger Rep_3 plasmid set could be classified (last column in Table 6). This is largely due to the predominance of R3-T1 (95 of 382 total) and R3-T2 (30 of 382) plasmids both of which were originally included in GR2 and encode, respectively, an Aci1 or Aci2 replication initiation protein. However, in the larger analysis, the R3-T3 (GR24) group, which is exemplified by pABTJ2 (36) is also abundant, including 45 members,.

**TABLE 7.** Rep_3 family types

| Type Rep | Plasmid name<sup>a</sup> | Aci | Number<br>95% (99%) | Group<sup>b</sup> | Protein id | Accession number |
| --- | --- | --- | --- | --- | --- | --- |
| R3-T1 | **pAB0057** (pA1-1)<sup>c</sup> | Aci1 | 95 (88) | GR2 | ACJ43223.1 | CP001183.2 |
| R3-T2 | pD72-1 (**pABV01**)<sup>d</sup> | Aci2 | 30 (29) | GR2/20 | AIH07953.1 | KM051986.1 |
| R3-T3 | pABTJ2 | - | 45 (44) | GR24 | AGG91013.1 | CP004359.1 |
| R3-T4 | **pMAC** (p2ATCC19606) | Aci9 | 15 (13) | GR8 | AAT09649.1 | AY541809.1 |
| R3-T5 | pABLAC1 | Aci3 | 13 (13) | GR3 | AIY39145.1 | CP007713.1 |
| R3-T6 | p1ZQ9 | - | 13 (13) | - | PQJ03469.1 | CM009083.3 |
| R3-T7 | **p3ABAYE** | - | 10 (7) | GR13 | CAM84695.1 | CU459140.1 |
| R3-T8 | **pAB02** | - | 10 (9) | GR12/29 | AAR00517.1 | AY228470.1 |
| R3-T9 | pABUH5-114 | - | 9 (9) | GR30 | WP_000095317.1 | NZ_AYOI01000002.1 |
| R3-T10 | **p2ABSDF** | - | 8 (8) | GR12 | CAP02944.1 | CU468232.1 |
| R3-T11 | p5ZQ3<sup>e</sup> | Aci4 | 8 (8) | GR4 | PQJ03708.1 | CM009651.1 |
| R3-T12 | p1ATCC19606 | - | 7 (6) | (GR2) | QFQ03441.1 | CP045108.1 |
| R3-T13 | pABA-6973 | - | 7 (5) | - | AUT39979.1 | CP026126.1 |
| R3-T14 | pAb242_9 | - | 5 (5) | (GR4)<sup>f</sup> | AUO31880.1 | KY984045.1 |
| R3-T15 | pABAY15001_6E | - | 5 (5) | - | QBN23345.1 | MK386684.1 |
| R3-T16 | pTG31986 | - | 5 (5) | - | QCD20890.1 | CP039342.1 |
| R3-T17 | pAba3207a | - | 5 (5) | GR26 | ANC38759.1 | CP015365.1 |
| R3-T18 | **p1ABAYE** | - | 4 (4) | GR11 | CAM84608.1 | CU459137.1 |
| R3-T19 | **pACICU1b** | Aci10 | 4 (4) | GR10 | QCS03997.1 | CP031381.2 |
| R3-T20 | pRCH51-3 | - | 4 (4) | - | AQT19035.1 | KY216144.1 |
| R3-T21 | pABA-2f10 | - | 4 (4) | - | AUT40247.1 | CP026129.1 |
| R3-T22 | pAb-MCR4.1 | - | 4 (4) | - | AYY91210.1 | CP033872.1 |
| R3-T23 | p1ZQ8 | - | 4 (3) | - | PQL85636.1 | CM009033.2 |
| R3-T24 | pK09-14 | - | 4 (4) | - | QER77251.1 | CP043954.1 |
| R3-T25 | pD36-4 | - | 3 (3) | GR31 | ALJ89824.1 | CP012956.1 |
| R3-T26 | pAba810CPa | - | 3 (3) | GR27 | AXG87071.1 | CP026340.1 |
| R3-T27 | pA52-2 | - | 3 (3) | - | QAB42494.1 | CP034094.1 |
| R3-T28 | pDETAB2 | - | 3 (3) | GR34 | QMS84089.1 | CP047975.1 |
| R3-T29 | p2Res13-Abat<sup>g</sup> | - | 3 (3) | - | QPF15404.1 | CP062922.1 |
| R3-T30 | pPM193665_5 | - | 2 (2) | - | QJG86394.1 | CP050420.1 |
| R3-T31 | pR2091 | Aci5<sup>h</sup> | 2 (0) | GR5 | CUW37058.1 | LN997847.1 |
| R3-T32 | pAb242_25<sup>i</sup> | Aci8<sup>i</sup> | 2 (2) | GR8/23 | AUO31913.1 | KY984047.1 |
| R3-T33 | p3FDAARGOS_540 | - | 2 (1) | - | AYX85221.1 | CP033751.1 |
| R3-T34 | pA1296_2 | - | 2 (2) | - | ATI40537.1 | CP018334.1 |
| R3-T35 | pNaval81-26 | - | 2 (2) | GR28 | WP_000185726.1 | NZ_AFDB02000003 |
| R3-T36 | pAb242_25 | - | 2 (2) | GR22 | AUO31910.1 | KY984047.1 |
| R3-T37 | **pAB1** | - | 1 | GR17 | ABO13860.1 | CP000522.1 |
| R3-T38 | **p1ABSDF** | - | 1 | GR1 | CAP02936.1 | CU468231.1 |
| R3-T39 | **p2ABSDF** | - | 1 | GR18 | CAP02966.1 | CU468232.1 |
| R3-T40 | **p3ABSDF** | - | 1 | GR7 | CAP02976.1 | CU468233.1 |
| R3-T41 | **p3ABSDF** | - | 1 | GR9 | CAP02983.1 | CU468233.1 |
| R3-T42 | **p3ABSDF** | - | 1 | GR15 | CAP02992.1 | CU468233.1 |
| R3-T43 | pRCH52-1 | (Aci7)<sup>j</sup> | 1 | GR3 | ALC76579.1 | KT346360.1 |
| R3-T44 | pFDAARGOS_540 | - | 1 | - | AYX85290.1 | CP033753.1 |
| R3-T45 | p29FS20-1 | - | 1 | - | QLF12435.1 | CP044520.1 |
| R3-T46 | pEH_gr13 | - | 1 | - | QBR82727.1 | CP038259.1 |
| R3-T47 | pE47_009 | - | 1 | - | _k | CP042565.1 |
| R3-T48 | p1Res13-Abat<sup>g</sup> | - | 1 | - | QPF15412.1 | CP062923.1 |
| R3-T49 | pABUH2a-5.6 | - | 1 | - | WP_032021082.1 | NZ_AYFZ01000080 |
| R3-T50 | pA1429b | - | 1 | - | QLB37617.1 | CP046900.1 |
| R3-T51 | p4ZQ2 | - | 1 | - | PQJ03811.1 | CM009050.2 |
| R3-T52 | pDT0544C | - | 1 | - | QLI38221.1 | CP053216.1 |
| R3-T53 | pDA33382-2-2 | - | 1 | - | AXB17573.1 | CP030107.1 |
| R3-T54 | pP7774 | - | 1 | - | QCR91187.1 | CP040262.1 |
| R3-T55 | pE47_008 | - | 1 | - | QFH47722.1 | CP042564.1 |
| R3-T56 | pVB2486_5 | - | 1 | - | QJH05174.1 | CP050408.1 |
| R3-T57 | pAba10324a | - | 1 | - | AXX46897.1 | CP023023.1 |
| R3-T58 | pA52-4 | - | 1 | - | QAB42527.1 | CP034096.1 |
| R3-T59 | pOCU_Ac16a_3 | - | 1 | - | _k | AP023080.1 |
| R3-T60 | p2ZQ2 | - | 1 | - | PQJ03830.1 | CM009647.1 |
| R3-T61 | pAC1633-4 | - | 1 | - | QOJ62393.1 | CP059302.1 |
| R3-T62 | pS30-1 | - | 1 | - | ARH59503.1 | KY617771.1 |
| R3-T63 | pD36-4 | - | 1 | - | ALJ89842.1 | CP012956.1 |
| R3-T64 | pDT01139C | - | 1 | - | QLI41769.1 | CP053221.1 |
| R3-T65 | pKCRI-43-1 | - | 1 | - | _k | LR026973.1 |
| R3-T66 | pE47_003 | - | 1 | - | QFH47692.1 | CP042559.1 |
| R3-T67 | pAC1633-2 | - | 1 | - | QOJ62400.1 | CP059303.1 |
| R3-T68 | pAb242_12 | - | 1 | GR21 | AUO31881.1 | KY984046.1 |
| R3-T69 | p135040 | - | 0<sup>l</sup> | GR19 | ACX70400.1 | GQ861437.1 |

<sup>a</sup> Names in bold are plasmids used in Bertini *et al.* 2010.
<sup>b</sup> groups assigned by Bertini *et al.*, 2010 (GR1-GR19), Lean (GR20), Cammeranesi *et al.*, 2020 (GR21-GR23), Salgado Camargo (GR24-GR33) and Liu *et al.* 2020 (GR34). Brackets indicate types falling within the GR guidelines at 74% *rep* gene identity.
<sup>c</sup> Bertini *et al.* 2010 included four Aci1 plasmids pAB2, pACICU1, p2ABAYE and pAB0057. pA1-1 included as the oldest known example.
<sup>d</sup> Plasmid pD72-1 replaces pABV01 used by Bertini *et al.* 2010, which is now known to be a hybrid of the majority of Aci1 and Aci2 sequences.
<sup>e</sup> partial sequence only in Bertini *et al.* 2010 (p844; GU978998).
<sup>f</sup> recorded as GR4 in Cammeranesi *et al.*, 2020 and is 90.79% identical to the GR4 reference.
<sup>g</sup> unnamed plasmids named here. Full strain names are p1Res13-Abat-PEA21-P4-01-A and p2Res13-Abat-PEA21-P4-01-A.
<sup>h</sup> representative *rep* is in a partial sequence only (p537; GU978999) in Bertini *et al.*, 2010.
<sup>i</sup> *rep* of pAb242\_25 (encoding AUO31913.1) is identical to a partial sequence (p11921; GU979000) corresponding to Aci8 in GR8 in Bertini. *et al.*, 2010. Also assigned Aci23 in Cammeranesi *et al.*, 2020.
<sup>j</sup> pRCH52-1 *rep* is 95.16% identical to *rep*Aci7 (p11921; GU978996) from a partial sequence. No complete plasmids with identical *rep* genes were detected.
<sup>k</sup> not available; sequence file not annotated.
<sup>l</sup> no complete plasmids correspond to a *rep* in a partial of p135040 (GQ861437) in Bertini. *et al.*, 2010.

In some cases, recombination has occurred between related types generating hybrid *rep* genes that complicate classification. Our detailed examination of the *rep* genes of a number of plasmids included in the same R3 type but that have <99% identity overall to the type representative revealed that they consisted of two or more segments, the largest segment with very high identity to the representative for the assigned type and the remainder closely matching another type representative. One example is the *rep* genes of pAB02 and pABIR from the original GR12 group where the pABIR *rep* differs from the *rep* in the remaining members of the group with differences clustered at the 3’-end. A further example is the *repAci1/rep*Aci2 hybrid found in pABVA01 (shown in Figure 2B), which encodes the Aci2 exemplar in the original classification. The phylogeny of the Rep proteins shown in Figure 3 illustrates how some types, for example R3-T21 and R3-T25 or R3-T2 and R3-T12, would amalgamate if the cut off used were reduced to 90% protein sequence identity.

**FIG 3.**
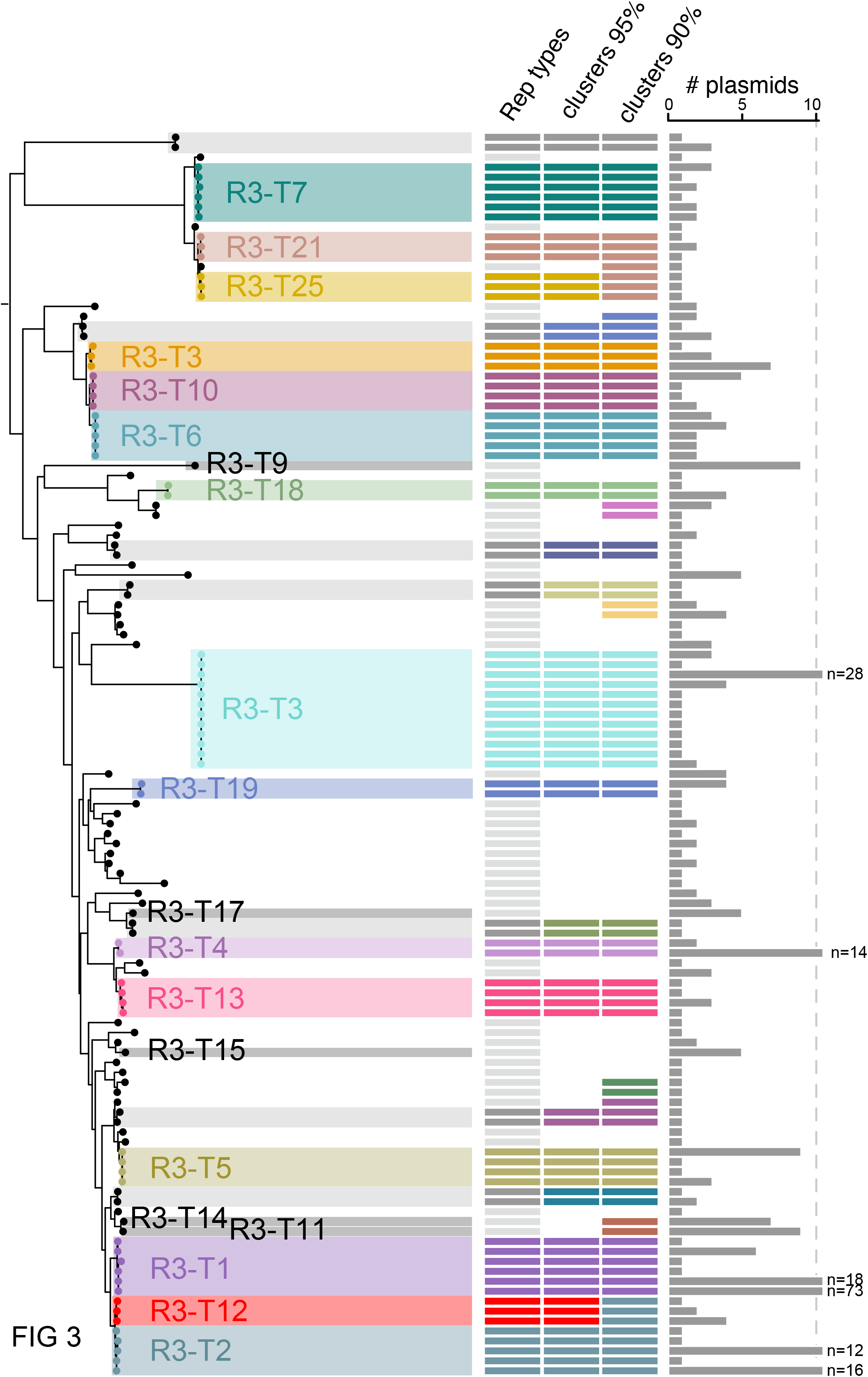
Maximum-likelihood phylogenetic relationship of replication initiation Rep sequences from the Rep_3 family. The tree comprises amino acid sequences encoded by n=125 unique nucleotide sequence variants from the Rep_3 family generated in IQ-TREE. The shading show either sequence variants assigned to the same Rep3 family (unlabelled) or those detected in at least n=5 plasmids (labelled). Columns are as follows: Rep types assigned based on a threshold of 95% nucleotide identity (those in light grey correspond to unique Rep types with a single nucleotide variant), clusters of Rep sequence variants grouped at 95% and 90% identity assigned with CD-HIT. The number of plasmids in which each Rep-encoding sequence variant was detected is shown on the right-hand side.

Plasmids encoding a Rep_3 family replication initiation protein are usually associated with iterons and use a theta replication mode. Indeed, iterons were identified in most of the Rep_3 encoding plasmids identified in the original classification (see Table 6). However, the presence of iterons was not systematically examined here.

### GR2 includes the R3-T1, R3-T2 and R3-T12 groups

An examination of the relationships between members of the R3-T1, R3-T2 and R3-T12 groups, which would all be included in the original GR2 group (2) provides insight into the reasons for increasing the cut-off to 95% identity. Most R3-T1 plasmids (encoding the Aci1 Rep protein) are found in multiply, extensively and pan resistant isolates belonging to global clones GC1 and GC2 which dominate the total genome sequences available currently. Indeed, the plasmid pA1-1 (GenBank accession number CP010782; (37)), which is found in the oldest GC1 isolate (1982) for which sequence data is available, is identical to pAB0057 (2004), indicating a long association of the plasmid with the clone. Many plasmids identical or nearly identical to pAB0057/pA1-1 were found among available complete sequences in a previous analysis (16). About half of the members of the R3-T1 group reported here are identical or nearly identical to pA1-1 with differences in recorded length generally due to failure to trim the overlap from linear contigs. The pAB5075/pA1-1 plasmid consists of a backbone and 2 *dif* modules (Figure 2A), though this was not initially recognised. Indeed, further R3-T1 plasmids carrying the same *rep*Aci1 gene have the same backbone but a different *dif* module in the first position or have a different set of *dif* modules, e.g. p2ABAYE and pD36-3 (Figure 2A). This lineage was also among the most abundant identified by others (10).

The *rep* gene encoding RepAci2 found in the R3-T2 type, also originally classified as GR2, is only approximately 80% identical to RepAci1 and was re-assigned to GR20 (5). However, here a third clearly distinct *rep* type (92.7% identity to *rep*Aci2) that would fall into the original GR2 group was found in p1ATCC19606 (38). This type was designated R3-T12. The close relationship between these three types can be seen in the phylogeny of R3 family Rep proteins (Figure 3).

### *Acinetobacter* Plasmid Typing database

To enable others to detect the presence of the *Acinetobacter* plasmid *rep* types defined here, we have made available databases comprising reference nucleotide sequences in GitHub (https://github.com/MehradHamidian/AcinetobacterPlasmidTyping). The databases comprise simple multi-fasta files that include a representative sequence for each *rep* type with information on the type plasmid (name and GenBank accession number) provided in the header. These files can be used to screen genome assemblies using BLASTN (run locally or using web-based services to screen read sets), or SRST2 ((39),https://github.com/katholt/srst2)using a threshold of 95% nucleotide identity to match the threshold used to define *rep* types. The databases can also be used to to screen sequencing reads for *rep* types, facilitating the detection of plasmid-derived contigs in short read assemblies for which no tool is currently available for *Acinetobacter*.

### Use of *Acinetobacter* Plasmid Typing database for read-based analysis

To demonstrate the use of the databases for typing, short read data sets used previously in an analysis of a collection of 41 GC1 isolates (40) were examined (see Methods). Analysis of the read data for the 36 isolates for which complete genomes were not available revealed that all but one of the isolates in the collection included a detectable plasmid *rep* gene (Table 8). Plasmids present in available complete genomes of five isolates (strain name shown in bold in Table 8) are also listed.

**TABLE 8.** Distribution of plasmid Rep types in GC1 isolates studied in Holt *et al*, 2016

| **Genomes<sup>a</sup>** | **Date** | **Country** | **ST** | **GenBank no.<sup>b</sup>** | **R3-T** | **RP-T** | **R1-T** |
| --- | --- | --- | --- | --- | --- | --- | --- |
| **A1** | 1982 | UK | 1 | CP010781-2 | T1<sup>c</sup> | - | - |
| **A388** | 2002 | Greece | 1 | CP024418-9 | T1, T19<sup>d</sup> | - | - |
| **AB0057** | 2004 | USA | 1 | CP001182-3 | T1<sup>e</sup> | - | - |
| **A85** | 2003 | Australia | 1 | CP021782-7 | T1<sup>f</sup> | T1 | T1, T2 |
| A297 | 1984 | Netherlands | 1 | FBWR | T1 | - | - |
| J1 | 1995 | Australia | 1 | FBWQ | T1 | T1 | - |
| J5 | 1997 | Australia | 1 | FBWP | T1 | T1 | - |
| WM98 | 1998 | Australia | 1 | UCPN | T1 | - | - |
| J7 | 1998 | Australia | 1 | FBWT | T1 | - | - |
| J10 | 1999 | Australia | 1 | FBWS | T1, T2 | - | - |
| D3208 | 1997 | Australia | 1 | FBWZ | T1 | - | - |
| D2 | 2006 | Australia | 1 | FBWY | T1, T15 | - | - |
| D62 | 2010 | Australia | 1 | FBWW | T1 | - | - |
| D30 | 2008 | Australia | 1 | FBXG | T1 | - | - |
| A83 | 2002 | Australia | 1 | FBWU | T1 | - | - |
| A92 | 2005 | Australia | 1 | FBWV | T1 | - | - |
| 6772166 | 2002 | Australia | 1 | FBWX | T1 | - | T2 |
| RBH3 | 2002 | Australia | 1 | FBXD | T1 | - | T2 |
| D13 | 2009 | Australia | 1 | FBXI | T14 | - | - |
| D15 | 2009 | Australia | 1 | FBXJ | T14 | - | - |
| G7 | 2003 | Australia | 1 | FBXF | T1 | T1 | - |
| D81 | 2010 | Australia | 1 | FBXC | T19 | T1 | - |
| D78 | 2010 | Australia | 1 | FBXH | T19 | T1 | - |
| Naval-83 | 2006 | USA | 20 | AMFK | - | T1 | - |
| AB058 | 2003 | USA | 20 | ADHA | T1, T8 | - | - |
| NIPH 527 | 1984 | Netherlands | 1 | APQW | T1 | - | - |
| NIPH 290 | 1994 | Netherlands | 1 | APRD | T9 | - | - |
| AB056 | 2004 | USA | 1 | ADGZ | T1 | - | T2 |
| AB059 | 2004 | USA | 1 | ADHB | T1 | T1 | - |
| 908-13 | 2007 | USA | 1 | AMHW | - | - | - |
| 909-02-7 | 2007 | USA | 1 | AMHZ | T1 | - | - |
| Canada-BC1 | 2007 | Canada | 1 | AMSZ | - | - | - |
| Canada-BC5 | 2007 | Canada | 1 | AFDN | T1 | - | - |
| IS-58 | 2008 | nr | 1 | AMGH | - | T1 | - |
| ANC 4097 | 2011 | nr | 1 | APRF | T1, T12, T57 | T1 | T4 |
| **D36** | 2008 | Australia | 81 | CP012952-6 | T1, T25, T63<sup>g</sup> | - | - |
| 6013113 | 2007 | UK | 81 | ACYR | T25, T63 | - | - |
| 6013150 | 2007 | UK | 81 | ACYQ | T1, T4, T25, T63 | - | - |
| OIFC074 | 2003 | USA | 19 | MADE | T1 | - | - |
| Naval-21 | 2006 | USA | 19 | AMSY | - | T1 | - |
| TG19582 | nr | nr | 1 | AMIV | T63 | - | - |

<sup>a</sup> Complete genomes are shown using bold letters.<sup>b</sup> All draft genome accession numbers include “00000000”.<sup>c</sup> encoded by pA1-1 (GenBank accession number CP010782).<sup>d</sup> Both R3-T1 and R3-T19 Reps are encoded by pA388 (GenBank accession number CP024419).<sup>e</sup> encoded by pAB0057 (GenBank accession number CP001183).<sup>f</sup> A85 carries 5 plasmids, pA85-1 (R1-T1), pA85-1a (R1-T2), pA85-1b (does not encode a Rep), pA85-2 (R3-T1) and pA85-3 (RP-T1).<sup>g</sup> D36 carries 4 plasmids. pD36-1 and pD36-2 (pRAY\*) (GenBank accession numbers CP012953.1 and CP012954.1, respectively) do not encode a Rep. pD36-3 (GenBank accession number CP012955.1) encodes a R3-T1 Rep and pD36-4 (GenBank accession number CP012956.1) encodes two Reps (R3-T25 and R3-T63).

An R3-T1 plasmid was detected in most isolates, including all five with a completed genome. Inspection of the relevant contigs revealed that in all isolates belonging to GC1 lineage 1, this plasmid was identical or nearly identical to pAB0057/pA1-1. However, in the ST81 group in lineage 2 (isolates D36, 6013113 and 6013150) the R3-T1 plasmid matched pD36-3 which includes different *dif* modules. Plasmid types R3-T25 and R3-T63 were also found in ST81 isolates 6013113 and 6013150 consistent with their presence in the complete genome of D36, as reported previously (17).

A R3-T19 (Aci10) type plasmid was detected in isolates D78 and D81 together with an RP-T1 (Aci6) plasmid. The RP-T1 plasmids were previously reported to be present and identical to pAb-G7-2 (40) and this was verified using a complete genome assembly of D78 (C.J. Harmer and R.M. Hall, unpublished). The R3-T19 plasmid in D78 was found to include the *oxa58* carbapenem resistance gene. Similarly, the presence of an R3-T14 type in the related isolates D13 and D15 was confirmed using a complete genome assembly for D13 (C.J. Harmer and R.M. Hall, unpublished). However, the complete D13 genome assembly included a further Rep_3 family plasmid that was of a type not present in the current database, indicating that updates to the plasmid typing scheme will be needed in the future.

## DISCUSSION

The analyses described here enabled development of a simple classification scheme which covers the plasmids found in *A. baumannii* that include a *rep* gene encoding an identifiable replication initiation or Rep protein. The cut-off of 95% nucleotide identity was chosen because it is readily able to accommodate minor sequence difference arising from evolution or from sequencing errors and also facilitates grouping of hybrid genes with the one type from which the majority of its sequence has been derived. The effect of raising the cut off is clear in Figure 3 where groups of three coherent types can be found within the former GR2 (R3-T1, R3-T2 and R3-T12) and within the former GR12 (R3-T6, R3-T8 and R3-T10).

The simple strategy applied here provides a framework for the research community to classify *Acinetobacter baumannii* plasmids as the simple rules will allow new plasmid types to be added to the scheme as they are discovered. Ultimately, this may include some of the known large conjugative plasmid families which likely encode a Rep protein of a type that has not yet been identified and verified experimentally and hence has not been assigned a Pfam. Once a potential *rep* gene is identified, these can be simply added into the scheme. This scheme should also be applicable to plasmids found in other *Acinetobacter* species as plasmid sharing is known to occur (14).

Because there is little variation between the *rep* genes in members of individual types in the R1 and RP groups and significant variation between the *rep* gene associated with each type, identification of additional types should be straightforward. Indeed, complete sequence of plasmids belonging to each R1 type have so far been identical. In the case of the RP-T1 and RP-T2 groups, members share very closely related backbones. However, in the case of the more diverse R3 group, the effect of recombination between moderately closely related plasmids may lead to complications in the future. One prime example is the hybrid *rep* genes that arise from recombination of two or more types potentially leading to formation of *rep* types that do not fit neatly into an existing category. Indeed, some examples of hybrids were identified here among plasmids with backbones of the iteron-*rep*-orfX configuration that include *dif* modules. However, a more detailed analysis of the more diverged members of the R3 types that include plasmids with *rep* genes that are less than 99% identical to the type representative will be needed to further explore these relationships. For this group, an improvement in the accuracy of annotation of *rep* genes is clearly needed. However, we note that in the RefSeq version of these plasmid sequences, the validated *rep* gene is annotated as *repM*, following Dorsey *et al*. (35) nomenclature, and orfX is annotated as a helix-turn-helix domain protein. The Aci classifications (*rep*Aci) that are applicable to some of the dominant types have been widely used and are likely to remain useful into the future.

The sequence files containing representative sequences of each type have been made available in a public GitHub repository and will simplify classification of competed plasmid genomes. These files will underpin the detection of plasmids in short read DNA sequence data, which is rarely undertaken currently due to the difficulties involved. This should lead to a more comprehensive approach to the surveillance of the role of plasmid-encoded functions in antibiotic resistance, virulence and environmental survival.

## MATERIALS AND METHODS

### Plasmid sequence data collection

Plasmids from *A. baumannii* whole genome sequence projects were downloaded from GenBank (https://www.ncbi.nlm.nih.gov/genbank/) in February 2021 and duplicate sequences from the same isolate were removed. Plasmids with sequences of poor quality were eliminated for various reasons such as sequencing errors, assembly issues, circularisation/trimming issues or the *rep* gene was incomplete. Reasons for removal are documented in Supplementary Table S1. After curation, there were 539 plasmid nucleotide sequences available from published genome projects.

To capture complete plasmids sequenced using traditional methods (not part of a genome project), the RefSeq (https://www.ncbi.nlm.nih.gov/refseq/) database was also searched using the search terms “Acinetobacter baumannii AND complete AND plasmid and srcdb_refseq[PROP]”.

All plasmids with accession numbers starting with CP, CM, CU etc. indicating they were from complete genomes were removed and the remainder curated as described above. This identified an additional 81 plasmids (including the 15 complete plasmids and 1 partial sequence described in (2) now completed, resulting in a final dataset comprising 621 complete plasmids. This data set was used to develop the typing scheme in this study.

Initially, plasmid entries with annotations were inspected manually, by searching for words such as replication or rep, to find any annotated *rep* genes. In the case of plasmids encoding a Rep_3 family Rep, the genes incorrectly annotated as *repA* were identified (see Results for details) and removed from the gene set. The remaining entries with no annotations or with no annotated *rep*/REP were initially further screened using low stringency BLASTn searches with each of the DNA sequences of *rep* genes that had been previously described in Bertini et al., (2). To find *rep* genes in the remainder of entries, they further examined using tBLASTn using the amino acid sequences of Rep proteins described in (2). In addition searches with the complete sequences or the backbone sequences of the plasmids previously found to be devoid of a *rep* gene including pRAY* (GenBank Accession number CP012954.1), pA85-1b (GenBank Accession number CP021785.1), pD36-1 (GenBank Accession number CP012953.1), pA297-3 (GenBank accession number), pNDM-BJ01 (GenBank accession number JQ001791.1) and pALWED1.1 (GenBank accession number CP082144.1) were used to identify related plasmids.

After identification of the plasmids encoding a Rep protein, the DNA sequences of *rep* genes identified using annotations, BLASTn and tBLASTn were extracted and used to create a local database used for further analysis of the *rep* genes. Amino acid sequence data for the Rep of each entry were extracted and used to populate a second local database used for further analysis.

### Clustering and phylogenetic analysis of the *rep*/Rep sequence data

Clusters comprising *rep* sequences at >80%, >85%, >90% and >95% nucleotide similarity were derived using CD-HIT Suite (https://github.com/weizhongli/cdhit) (41). To study the relationships of the Rep protein sequences, the nucleotide sequences for the extracted *rep* sequences were translated to amino acid sequences with EMBOSS Transeq and aligned with MUSCLE v3.8.31 (42). The aligned sequence file was used to infer a maximum likelihood phylogeny with IQ-TREE version 1.6.10 (43, 44) with the VT+F+G4 model (selected from the -m test model selection flag) with 100 bootstrap replicates. The final tree was visualised in FigTree v1.4.4 (http://tree.bio.ed.ac.uk/) and tree annotation generated with the plotTree code (github.com/katholt/plotTree) in R v1.1.456.

## SUPPLEMENTAL MATERIAL

Supplemental material is available online only.

**SUPPLEMENTAL FILE 1**, Supplemental Material, PDF file, 320 KB.

## ACKNOWLEDGEMENTS

This project and M.H. were supported by an Australian Research Council (ARC) DECRA fellowship (fellowship DE200100111).

## AUTHORS AND CONTRIBUTORS

Conceptualization: MH. Data curation M.C.L, J.K, MH. Formal Analysis: M.C.L, J.K, MH. Funding acquisition M.H. Investigation M.C.L, J.K, K.E.H, R.M.H, MH. Methodology M.C.L, J.K, K.E.H, R.M.H, MH. Visualization M.C.L, MH. Writing – original draft, R.M.H, MH. Writing – review & editing M.C.L, K.E.H, R.M.H, MH.

## REFERENCES

1. Carattoli A, Zankari E, García-Fernández A, Voldby Larsen M, Lund O, Villa L, Møller Aarestrup F, Hasman H. 2014. In silico detection and typing of plasmids using PlasmidFinder and plasmid multilocus sequence typing. Antimicrob Agents Chemother 58:3895–3903.

2. Bertini A, Poirel L, Mugnier PD, Villa L, Nordmann P, Carattoli A. 2010. Characterization and PCR-based replicon typing of resistance plasmids in Acinetobacter baumannii. Antimicrob Agents Chemother 54:4168–4177.

3. Towner KJ, Evans B, Villa L, Levi K, Hamouda A, Amyes SG, Carattoli A. 2011. Distribution of intrinsic plasmid replicase genes and their association with carbapenem-hydrolyzing class D B-lactamase genes in European clinical isolates of Acinetobacter baumannii. Antimicrob Agents Chemother 55:2154–2159.

4. Fu Y, Jiang J, Zhou H, Jiang Y, Fu Y, Yu Y, Zhou J. 2014. Characterization of a novel plasmid type and various genetic contexts of blaOXA-58 in Acinetobacter spp. from multiple cities in China. PLoS One 9:e84680.

5. Lean SS, Yeo CC. 2017. Small, enigmatic plasmids of the nosocomial pathogen, Acinetobacter baumannii: Good, bad, who knows? Front Microbiol 8:1547.

6. Liu L-L, Ji S-J, Ruan Z, Fu Y, Fu Y-Q, Wang Y-F, Yu Y-S. 2015. Dissemination of blaOXA-23 in Acinetobacter spp. in China: main roles of conjugative plasmid pAZJ221 and transposon Tn2009. Antimicrob Agents Chemother 59:1998–2005.

7. Blackwell GA, Hall RM. 2017. The tet39 determinant and the msrE-mphE genes in Acinetobacter plasmids are each part of discrete modules flanked by inversely oriented pdif (XerC-XerD) sites. Antimicrob Agents Chemother 61:e00780–17.

8. Hua X, Pan C, Zhu L, Liu Z, Xu Q, Wang H, Yu Y. 2017. Complete genome sequence of Acinetobacter baumannii A1296 (ST1469) with a small plasmid harbouring the tet(39) tetracycline resistance gene. J Glob Antimicrob Resist 1:105–107.

9. Cameranesi MM, Limansky AS, Moran-Barrio J, Repizo GD, Viale AM. 2017. Three novel Acinetobacter baumannii plasmid replicase-homology groups inferred from anaysis of a multidrug-resistant clinical strain isolated in Argentina. J Infect Dis Epidemiol 3:046.

10. Salgado-Camargo AD, Castro-Jaimes S, Gutierrez-Rios RM, Lozano LF, Altamirano-Pacheco L, Silva-Sanchez J, Pérez-Oseguera Á, Volkow P, Castillo-Ramírez S, Cevallos MA. 2020. Structure and evolution of Acinetobacter baumannii plasmids. Front Microbiol 11:1283.

11. Liu H, Moran RA, Chen Y, Doughty EL, Hua X, Jiang Y, Xu Q, Zhang L, Blair JMA, McNally A, van Schaik W, Yu Y. 2021. Transferable Acinetobacter baumannii plasmid pDETAB2 encodes OXA-58 and NDM-1 and represents a new class of antibiotic resistance plasmids. J Antimicrob Chemother 76:1130–1134.

12. Hamidian M, Ambrose SJ, Hall RM. 2016. A large conjugative Acinetobacter baumannii plasmid carrying the sul2 sulphonamide and strAB streptomycin resistance genes. Plasmid 87-88:43–50.

13. Hamidian M, Nigro SJ, Hall RM. 2012. Variants of the gentamicin and tobramycin resistance plasmid pRAY are widely distributed in Acinetobacter. J Antimicrob Chemother 67:2833–2836.

14. Mindlin S, Beletsky A, Rakitin A, Mardanov A, Petrova M. 2020. Acinetobacter plasmids: Diversity and development of classification strategies. Front Microbiol 11:588410.

15. Salto IP, Torres Tejerizo G, Wibberg D, Pühler A, Schlüter A, Pistorio M. 2018. Comparative genomic analysis of Acinetobacter spp. plasmids originating from clinical settings and environmental habitats. Sci Rep 8:7783.

16. Blackwell GA, Hall RM. 2019. Mobilisation of a small Acinetobacter plasmid carrying an oriT transfer origin by conjugative RepAci6 plasmids. Plasmid 102.

17. Hamidian M, Hall RM. 2018. Genetic structure of four plasmids found in Acinetobacter baumannii isolate D36 belonging to lineage 2 of global clone 1. PLoS One 13:e0204357.

18. Mindlin S, Beletsky A, Mardanov A, Petrova M. 2019. Adaptive dif modules in permafrost strains of Acinetobacter lwoffii and their distribution and abundance among present day Acinetobacter Strains. Front Microbiol 10:632.

19. Hua X, Xu Q, Zhou Z, Ji S, Yu Y. 2019. Relocation of Tn2009 and characterization of an ABGRI3-2 from re-sequenced genome sequence of Acinetobacter baumannii MDR-ZJ06. J Antimicrob Chemother 74:1153–1155.

20. Blackwell GA, Holt KE, Bentley SD, Hsu LY, Hall RM. 2017. Variants of AbGRI3 carrying the armA gene in extensively antibiotic-resistant Acinetobacter baumannii from Singapore. J Antimicrob Chemother 72:1031–1039.

21. Hamidian M, Hawkey J, Wick R, Holt, K.E., Hall RM. 2019. Evolution of a clade of Acinetobacter baumannii global clone 1, lineage 1 via acquisition of carbapenem and aminoglycoside resistance genes and dispersion of ISAba1. Microbial Genomics 5:mgen.0.000242.

22. Hu H, Hu Y, Pan Y, Liang H, Wang H, Wang X, Hao Q, Yang X, Yang X, Xiao X, Luan C, Yang Y, Cui Y, Yang R, Gao GF, Song Y, Zhu B. 2012. Novel plasmid and its variant harboring both a *bla*(NDM-1) gene and type IV secretion system in clinical isolates of Acinetobacter lwoffii. Antimicrob Agents Chemother 56:1698–1702.

23. Mindlin S, Maslova O, Beletsky A, Nurmukanova V, Zong Z, Mardanov A, Petrova M. 2021. Ubiquitous conjugative mega-plasmids of Acinetobacter species and their role in horizontal transfer of multi-drug resistance. Front Microbiol 12:728644.

24. Nigro SJ, Hall RM. 2011. GIsul2, a genomic island carrying the sul2 sulphonamide resistance gene and the small mobile element CR2 found in the Enterobacter cloacae subspecies cloacae type strain ATCC 13047 from 1890, Shigella flexneri ATCC 700930 from 1954 and Acinetobacter baumannii ATCC 17978 from 1951. J Antimicrob Chemother 66:2175–2176.

25. Harmer CJ, Lebreton F, Stam J, McGann PT, Hall RM. 2022. Complete genome of the extensively antibiotic-resistant GC1 Acinetobacter baumannii isolate MRSN 56 reveals a novel route to fluoroquinolone resistance. J Antimicrob Chemother 77:1851–1855.

26. Wick RR, Judd LM, Wyres KL, Holt KE. 2021. Recovery of small plasmid sequences via Oxford Nanopore sequencing. Microb Genom 7:0.00631.

27. Hamidian M, Hall RM. 2014. pACICU2 is a conjugative plasmid of Acinetobacter carrying the aminoglycoside resistance transposon TnaphA6. J Antimicrob Chemother 69:1146–1148.

28. Hamidian M, Holt KE, Pickard D, Dougan G, Hall RM. 2014. A GC1 Acinetobacter baumannii isolate carrying AbaR3 and the aminoglycoside resistance transposon TnaphA6 in a conjugative plasmid. J Antimicrob Chemother 69:955–958.

29. Hamidian M, Kenyon JJ, Holt KE, Pickard D, Hall RM. 2014. A conjugative plasmid carrying the carbapenem resistance gene *bla*_OXA-23_ in AbaR4 in an extensively resistant GC1 Acinetobacter baumannii isolate. J Antimicrob Chemother 69:2625–2628.

30. Gallagher LA, Ramage E, Weiss EJ, Radey M, Hayden HS, Held KG, Huse HK, Zurawski DV, Brittnacher MJ, Manoil C. 2015. Resources for genetic and genomic analysis of emerging pathogen Acinetobacter baumannii. J Bacteriol 15;197:2027–2035.

31. Nigro SJ, Holt KE, Pickard D, Hall RM. 2015. Carbapenem and amikacin resistance on a large conjugative Acinetobacter baumannii plasmid. J Antimicrob Chemother 70:1259-1261.

32. Huang H, Yang Z-L, Wu X-M, Wang Y, Liu Y-J, Luo H, Lv X, Gan Y-R, Song S-D, Gao F. 2012. Complete genome sequence of Acinetobacter baumannii MDR-TJ and insights into its mechanism of antibiotic resistance. J Antimicrob Chemother 67:2825–32.

33. Liu CC, Kuo HY, Tang CY, Chang KC, Liou ML. 2014. Prevalence and mapping of a plasmid encoding a type IV secretion system in Acinetobacter baumannii. Genomics 104:215–223.

34. D’Andrea MM, Giani T, D’Arezzo S, Capone A, Petrosillo N, Visca P, Luzzaro F, Rossolini GM. 2009. Characterization of pABVA01, a plasmid encoding the OXA-24 carbapenemase from Italian isolates of Acinetobacter baumannii. Antimicrob Agents Chemother 53:3528-3233.

35. Dorsey CW, Tomaras AP, Actis LA. 2006. Sequence and organization of pMAC, an Acinetobacter baumannii plasmid harboring genes involved in organic peroxide resistance. Plasmid 56:112–123.

36. Huang H, Dong Y, Yang ZL, Luo H, Zhang X, Gao F. 2014. Complete sequence of pABTJ2, a plasmid from Acinetobacter baumannii MDR-TJ, carrying many phage-like elements. Genomics Proteomics Bioinformatics 12:172–177.

37. Holt KE, Hamidian M, Kenyon JJ, Wynn MT, Hawkey J, D. P, Hall RM. 2015. Genome sequence of Acinetobacter baumannii strain A1, an early example of antibiotic-resistant Global Clone 1. Genome Announc 3:e00032–15.

38. Hamidian M, Blasco L, Tillman LN, To J, Tomas M, Myers GSA. 2020. Analysis of complete genome sequence of Acinetobacter baumannii strain ATCC 19606 reveals novel mobile genetic elements and novel prophage. Microorganisms 8:1851.

39. Inouye M, Conway TC, Zobel J, Holt KE. 2012. Short read sequence typing (SRST): multilocus sequence types from short reads. BMC Genomics 13:338.

40. Holt KE, Kenyon JJ, Hamidian M, Schultz MB, Pickard DJ, Dougan G, Hall RM. 2016. Five decades of genome evolution in the globally distributed, extensively antibiotic-resistant Acinetobacter baumannii global clone 1. Microb Genomics 2:mgen.0.000052.

41. Li W, Godzik A. 2006. Cd-hit: a fast program for clustering and comparing large sets of protein or nucleotide sequences. Bioinformatics 22:1658–1659.

42. Edgar RC. 2004. MUSCLE: multiple sequence alignment with high accuracy and high throughput. Nucleic Acids Res 32:1792–1797.

43. Kalyaanamoorthy S, Minh BQ, Wong TKF, von Haeseler A, Jermiin LS. 2017. ModelFinder: fast model selection for accurate phylogenetic estimates. Nature Methods 14:587–589.

44. Nguyen L-T, Schmidt H, A., von Haeseler A, Minh BQ. 2015. IQ-TREE: A fast and effective stochastic algorithm for estimating maximum-likelihood phylogenies. Molec Biol Evolution, 32:268–274.

