## Supplemental Table 1, 2 and 3 for "Detection and typing of plasmids in *Acinetobacter baumannii* using *rep* genes encoding replication initiation proteins"

**Table S1** Plasmids excluded from study

| Accession number | Reason for exclusion |
|---|---|
| CP050525.1 | Not plasmid sequence – RNA |
| CP040042.1 | Not plasmid sequence – Chromosome |
| CP040041.1 | Not plasmid sequence – Chromosome |
| CP040048.1 | Not plasmid sequence – Chromosome |
| CP020583.1 | Contaminated sample |
| CP016296 | Duplicated accession number |
| CP016297 | Duplicated accession number |
| CP016299 | Duplicated accession number |
| CP016301 | Duplicated accession number |
| CP016302 | Duplicated accession number |
| CP017655 | Duplicated accession number |
| CP047974 | Duplicated accession number |
| CP047975 | Duplicated accession number |
| CP047976 | Duplicated accession number |
| KP890934.1 | Assembly issue - Frameshift |
| CP035931.1 | Assembly issue - Frameshift |
| CP033517.1 | Assembly issue - Frameshift |
| NZ_AFDA02000007.1 | Assembly issue - Frameshift |
| CP030084.1 | Assembly issue - Frameshift |
| CP032741.1 | Assembly issue - Frameshift |
| HG977528.1 | Assembly issue - Frameshift |
| CP033519.1 | Assembly issue - Frameshift |
| NZ_AFCZ02000004.1 | Assembly issue - Frameshift |
| CP035674.1 | Assembly issue - Frameshift |
| CP035675.1 | Assembly issue - Frameshift |
| CP040046.1 | Assembly issue - Frameshift |
| CP033518.1 | Assembly issue - Frameshift |
| CP040081.1 | Assembly issue - Frameshift |
| CP061542.1 | Assembly issue - Frameshift |
| CP020575.1 | Assembly issue - Frameshift |
| CP040082.1 | Assembly issue - Frameshift |
| CP018862.2 | Assembly issue - Gap |
| CP065393.1 | Assembly issue - Gap |
| CP032216.1 | Incomplete plasmid sequence |
| CP032219.1 | Incomplete plasmid sequence |
| AP022837.1 | Reassembled/ resubmitted plasmid |
| AP022838.1 | Reassembled/ resubmitted plasmid |
| CP000865.1 | Reassembled/ resubmitted plasmid |
| CP019218.1 | Reassembled/ resubmitted plasmid |
| CP035052.1 | Reassembled/ resubmitted plasmid |
| CP039024.1 | Reassembled/ resubmitted plasmid |
| CP039026.1 | Reassembled/ resubmitted plasmid |
| CP039027.1 | Reassembled/ resubmitted plasmid |
| CP039029.1 | Reassembled/ resubmitted plasmid |
| CP039030.1 | Reassembled/ resubmitted plasmid |
| CP046655.1 | Reassembled/ resubmitted plasmid |
| CP049364.1 | Reassembled/ resubmitted plasmid |
| CP049365.1 | Reassembled/ resubmitted plasmid |
| CP050398.1 | Reassembled/ resubmitted plasmid |
| CP053099.1 | Reassembled/ resubmitted plasmid |
| CP053100.1 | Reassembled/ resubmitted plasmid |
| CP059042.1 | Reassembled/ resubmitted plasmid |

| | |
|---|---|
| JQ904627.1 | Reassembled/ resubmitted plasmid |
| KJ493819.2 | Reassembled/ resubmitted plasmid |
| MK360916.1 | Reassembled/ resubmitted plasmid |
| CP000864.1 | Reassembled/ resubmitted plasmid |
| CP015485.1 | Assembly issue - incomplete |
| CP050399.1 | Plasmid segment |
| CP050396.1 | Plasmid segment |
| CP050397.1 | Plasmid segment |
| MK134375.1 | Assembly issue – shortened *rep* |
| CP042995.1 | Wrong species |
| CP042996.1 | Wrong species |
| CP045561.1 | Wrong species |
| CM009040.2 | Assembly issue – Chromosomal contig |
| CM009044.2 | Assembly issue – Chromosomal contig |
| CM009649.1 | Assembly issue – Chromosomal contig |
| CP003907.1 | Circularised genomic island |
| CP003908.1 | Circularised genomic island |
| CM008887.1 | Circularised genomic island |
| CP007579.1 | Circularised genomic island |
| CP038646.1 | Circularised genomic island |
| HG977524.1 | Assembly issue – shortened *rep* |
| CP027184.1 | Assembly issue – shortened *rep* |
| CP017643.1 | Assembly issue – shortened *rep* |
| CM008887.2 | Assembly issue – shortened *rep* |

**Table S2** Properties of Rep_PriCT plasmids

| Acession number | Biosample | strain_name | Plasmid name | Length(bp) | Rep_family | Type | Bertini scheme | Locus/Protein_id |
|---|---|---|---|---|---|---|---|---|
| AEOY01000095.1 | SAMN02471598 | 3990 | p1ABST2 | 63320 | Rep | RP-T1 | Aci6 | - |
| AFDB02000004.1 | SAMN00114928 | Naval-81 | pNaval81-67 | 67012 | Rep | RP-T1 | Aci6 | EJP56764.1 |
| AFDL01000006.1 | SAMN02436631 | OIFC143 | pOIFC143-70 | 69518 | Rep | RP-T1 | Aci6 | EJG16496.1 |
| ALII01000020.1 | SAMN02436485 | IS-123 | pIS123-67 | 67025 | Rep | RP-T1 | Aci6 | EJO37624.1 |
| AP023078.1 | SAMD00059694 | OCU_Ac16a | pOCU_Ac16a_1 | 73028 | Rep | RP-T1 | Aci6 | - |
| AYOH01000010.2 | SAMN02597375 | UH9907 | pABUH1-74 | 74089 | Rep | RP-T1 | Aci6 | ETR49020.1 |
| CM009084.2 | SAMN08043814 | ZQ1 | p2ZQ9 | 62339 | Rep | RP-T1 | Aci6 | PLK14912.1 |
| CM009650.1 | SAMN08093362 | ZQ3 | p4ZQ3 | 68115 | Rep | RP-T1 | Aci6 | PQJ03752.1 |
| CP001922.1 | SAMN02603494 | 1656-2 | ABKp1 | 74451 | Rep | RP-T1 | Aci6 | ADX05416.1 |
| CP002524.1 | SAMN02603891 | TCDC-AB0715 | p2ABTCDC0715 | 70894 | Rep | RP-T1 | Aci6 | ADX94329.1 |
| CP007580.1 | SAMN02471421 | AC30 | pAC30c | 71433 | Rep | RP-T1 | Aci6 | AHX67284.1 |
| CP008707.1 | SAMN02894434 | AB5075-UW | p1AB5075 | 83610 | Rep | RP-T1 | Aci6 | AKA33572.1 |
| CP008851.1 | SAMN02714232 | AC29 | pAC29b | 74749 | Rep | RP-T1 | Aci6 | AKB09352.1 |
| CP012008.1 | SAMN03817045 | Ab04-mff | pAB04-2 | 87569 | Rep | RP-T1 | Aci6 | AKQ32652.1 |
| CP014216.1 | SAMN04448797 | YU-R612 | p2YU-R612 | 74241 | Rep | RP-T1 | Aci6 | AMC17795.1 |
| CP014292.1 | SAMN04457967 | AB34299 | p2AB34299 | 84967 | Rep | RP-T1 | Aci6 | AQU58831.1 |
| CP015366.1 | SAMN04485290 | 3207 | pAba3207b | 80546 | Rep | RP-T1 | Aci6 | ANC38839.1 |
| CP016297.1 | SAMN04096368 | CMC-CR-MDR-Ab4 | pCMCVTAb2-Ab4 | 74090 | Rep | RP-T1 | Aci6 | APQ87400.1 |
| CP016302.1 | SAMN04096370 | CMC-CR-MDR-Ab66 | pCMCVTAb2-Ab66 | 73188 | Rep | RP-T1 | Aci6 | APQ95054.1 |
| CP017645.1 | SAMN05601667 | KAB02 | pKAB02 | 81227 | Rep | RP-T1 | Aci6 | AOX75640.1 |
| CP017647.1 | SAMN05601668 | KAB03 | pKAB03 | 73891 | Rep | RP-T1 | Aci6 | AOX79427.1 |
| CP017649.1 | SAMN05601669 | KAB04 | pKAB04 | 121485 | Rep | RP-T1 | Aci6 | AOX83318.1 |
| CP017651.1 | SAMN05601670 | KAB05 | pKAB05 | 70884 | Rep | RP-T1 | Aci6 | AOX87285.1 |
| CP017653.1 | SAMN05601671 | KAB06 | pKAB06 | 70884 | Rep | RP-T1 | Aci6 | AOX91169.1 |
| CP017655.1 | SAMN05601672 | KAB07 | pKAB07 | 72073 | Rep | RP-T1 | Aci6 | AOX94964.1 |
| CP017657.1 | SAMN05601673 | KAB08 | pKAB08 | 101406 | Rep | RP-T1 | Aci6 | AOX98817.1 |
| CP019115.1 | SAMN06211572 | MDR-CQ | pMDR-CQ | 75390 | Rep | RP-T1 | Aci6 | QDR95729.1 |
| CP020573.1 | SAMN06650264 | 15A5 | p15A5_1 | 74241 | Rep | RP-T1 | Aci6 | ARF91039.1 |
| CP020577.1 | SAMN06650263 | SSA12 | pSSA12_1 | 73264 | Rep | RP-T1 | Aci6 | ARF94730.1 |
| CP020580.1 | SAMN06650261 | SSMA17 | pSSMA17_1 | 90973 | Rep | RP-T1 | Aci6 | ARG03955.1 |
| CP020582.1 | SAMN06650260 | JBA13 | pJBA13_1 | 90972 | Rep | RP-T1 | Aci6 | ARG07627.1 |
| CP020589.1 | SAMN06650243 | 15A34 | p15A34_1 | 72076 | Rep | RP-T1 | Aci6 | ARG18954.1 |
| CP020593.1 | SAMN06650241 | USA2 | pUSA2_1 | 74241 | Rep | RP-T1 | Aci6 | ARG29677.1 |
| CP020594.1 | SAMN06650240 | USA15 | pUSA15_1 | 98301 | Rep | RP-T1 | Aci6 | ARG29717.1 |
| CP021322.1 | SAMN07135563 | XH731 | pXH731 | 64557 | Rep | RP-T1 | Aci6 | AWW87056.1 |
| CP021787.1 | SAMN07125723 | A85 | pA85-3 | 86334 | Rep | RP-T1 | Aci6 | ASF79226.1 |
| CP023028.1 | SAMN07520233 | 10042 | pAba10042b | 110728 | Rep | RP-T1 | Aci6 | AXX50773.1 |
| CP023030.1 | SAMN07520232 | 9102 | pAba9102a | 95206 | Rep | RP-T1 | Aci6 | AXX54541.1 |
| CP023033.1 | SAMN07284119 | 7847 | pAba7847b | 80546 | Rep | RP-T1 | Aci6 | AXW92641.1 |
| CP024578.1 | SAMN07945345 | AbPK1 | pAbPK1b | 79335 | Rep | RP-T1 | Aci6 | ATR89566.1 |
| CP025267.1 | SAMN08054861 | SMC_Paed_Ab_BL01 | pSMC_AB_BL01_1 | 74241 | Rep | RP-T1 | Aci6 | AUG14723.1 |
| CP026706.1 | SAMN04014897 | AR_0056 | tig00000059_pilon | 72105 | Rep | RP-T1 | Aci6 | AVE44386.1 |
| CP026712.1 | SAMN04014904 | AR_0063 | unitig_2_pilon | 72990 | Rep | RP-T1 | Aci6 | AVE56836.1 |
| CP026946.1 | SAMN07977762 | S1 | pAbS1_03 | 72204 | Rep | RP-T1 | Aci6 | AVG28625.1 |
| CP027121.1 | SAMN04014897 | AR_0056 | p2AR_0056 | 72105 | Rep | RP-T1 | Aci6 | AVN03791.1 |
| CP027243.1 | SAMN08364584 | WCHAB005078 | p1_005078 | 70796 | Rep | RP-T1 | Aci6 | AVN12622.1 |
| CP029571.1 | SAMN09241862 | DA33098 | pDA33098-71 | 71234 | Rep | RP-T1 | Aci6 | AWO18521.1 |
| CP030109.1 | SAMN09460321 | DA33382 | pDA33382-85 | 84678 | Rep | RP-T1 | Aci6 | AXB17624.1 |
| CP031382.1 | SAMN09302593 | ACICU | pACICU2 | 70101 | Rep | RP-T1 | Aci6 | QCS04021.1 |
| CP033245.1 | SAMN07520235 | 7835 | pAba7835b | 80584 | Rep | RP-T1 | Aci6 | QFY70971.1 |
| CP035047.1 | SAMN05238672 | ABUH793 | p74.1Kbp | 74091 | Rep | RP-T1 | Aci6 | QAS99722.1 |
| CP036286.1 | SAMN10261590 | TG60155 | p60155_3 | 71234 | Rep | RP-T1 | Aci6 | QBH55839.1 |
| CP038264.1 | SAMN10386510 | LEV1449/17Ec | pEC_gr6 | 76956 | Rep | RP-T1 | Aci6 | QBR79239.1 |
| CP039994.1 | SAMN10261544 | TG22182 | pTG22182_1 | 71233 | Rep | RP-T1 | Aci6 | QCO84474.1 |
| CP040043.1 | SAMN11554497 | VB958 | p3VB958 | 82500 | Rep | RP-T1 | Aci6 | QCP18569.1 |
| CP042842.1 | SAMN12399660 | ATCC BAA-1790 | pATCCBAA-1790 | 67023 | Rep | RP-T1 | Aci6 | QEE59151.1 |
| CP050392.1 | SAMN14410107 | VB11737 | pVB11737_1 | 70712 | Rep | RP-T1 | Aci6 | QJH08823.1 |
| CP050404.1 | SAMN14414761 | VB2486 | pVB2486_1 | 99090 | Rep | RP-T1 | Aci6 | QJH05026.1 |
| CP050413.1 | SAMN14420253 | PM192696 | pPM192696_1 | 70098 | Rep | RP-T1 | Aci6 | QJG90047.1 |
| CP050909.1 | SAMN14308866 | DT-Ab022 | p2DT-Ab022 | 71194 | Rep | RP-T1 | Aci6 | QIX36610.1 |
| CP050912.1 | SAMN14308864 | DT-Ab020 | p2DT-Ab020 | 71194 | Rep | RP-T1 | Aci6 | QIX40477.1 |
| CP050915.1 | SAMN14308853 | DT-Ab007 | pDT-Ab007 | 92021 | Rep | RP-T1 | Aci6 | QIX44206.1 |
| CP050917.1 | SAMN14308849 | DT-Ab003 | p2DT-Ab003 | 71194 | Rep | RP-T1 | Aci6 | QIX48008.1 |
| CP061516.1 | SAMN12391854 | CFSAN093710 | pCFSAN093710_2 | 83626 | Rep | RP-T1 | Aci6 | QNV20001.1 |
| CP062921.1 | SAMN16304032 | Res13-Abat-PEA21-P4-01-A | p3Res13-Abat | 72831 | Rep | RP-T1 | Aci6 | QPF15346.1 |
| CP066230.1 | SAMN17073618 | G20AB011 | pG20AB011-1 | 73339 | Rep | RP-T1 | Aci6 | QQD98159.1 |
| CP066233.1 | SAMN17073617 | G20AB010 | pG20AB010-1 | 73360 | Rep | RP-T1 | Aci6 | QQD94357.1 |
| CP066236.1 | SAMN17073616 | G20AB009 | pG20AB009-1 | 73361 | Rep | RP-T1 | Aci6 | QQE05740.1 |
| CP066238.1 | SAMN17073615 | G20AB007 | pG20AB007-1 | 73361 | Rep | RP-T1 | Aci6 | QQE01989.1 |
| CP069841.1 | SAMN16357502 | FDAARGOS_1360 | p2FDAARGOS_1360 | 76933 | Rep | RP-T1 | Aci6 | QRR53484.1 |
| CP072124.1 | SAMN18396008 | KSK1 | p2KSK1 | 87529 | Rep | RP-T1 | Aci6 | QTH58400.1 |
| CP072272.1 | SAMN18452234 | KSK6 | p2KSK6 | 68224 | Rep | RP-T1 | Aci6 | QTK45694.1 |
| CP072277.1 | SAMN18452334 | KSK7 | p2KSK7 | 68225 | Rep | RP-T1 | Aci6 | QTK62065.1 |
| CP072282.1 | SAMN18452342 | KSK10 | p2KSK10 | 87529 | Rep | RP-T1 | Aci6 | QTK53937.1 |
| CP072287.1 | SAMN18452659 | KSK11 | p2KSK11 | 87529 | Rep | RP-T1 | Aci6 | QTK58013.1 |
| CP072292.1 | SAMN18452698 | KSK18 | p2KSK18 | 68225 | Rep | RP-T1 | Aci6 | QTK66135.1 |
| CP072297.1 | SAMN18452699 | KSK19 | p2KSK19 | 87529 | Rep | RP-T1 | Aci6 | QTK70246.1 |
| CP072302.1 | SAMN18452712 | KSK20 | p2KSK20 | 68309 | Rep | RP-T1 | Aci6 | QTK74363.1 |
| CP072400.1 | SAMN18451305 | KSK2 | p2KSK2 | 87529 | Rep | RP-T1 | Aci6 | QTL10322.1 |
| HG977523.1 | SAMEA3158456 | CS01 | pCS01A | 63720 | Rep | RP-T1 | Aci6 | - |
| HG977527.1 | SAMEA3158506 | CR17 | pCR17A | 63795 | Rep | RP-T1 | Aci6 | - |
| KF669606.1 | SAMN14225999 | G7 | pAb-G7-2 | 70100 | Rep | RP-T1 | Aci6 | AGY56182.1 |
| KF889012.1 | SAMN02603667 | TYTH-1 | pAB_CC | 65890 | Rep | RP-T1 | Aci6 | AII26481.1 |
| KM051846.1 | SAMN14226465 | D72 | pD72-2 | 70102 | Rep | RP-T1 | Aci6 | AIH07963.1 |
| KM977710.1 | SAMN14226502 | D46 | pD46-3 | 74916 | Rep | RP-T1 | Aci6 | AIZ49259.1 |
| KR535992.1 | SAMN14226627 | A105 | pA105-1 | 70098 | Rep | RP-T1 | Aci6 | ALN43326.1 |
| KU549175.1 | SAMN14227264 | C13 | pC13-2 | 103871 | Rep | RP-T1 | Aci6 | AMQ95288.1 |
| KX230794.1 | SAMN14227145 | MAL | pMAL-2 | 67025 | Rep | RP-T1 | Aci6 | ANR95813.1 |
| KY022424.1 | SAMN14227393 | Ab8098 | pAb8098 | 82667 | Rep | RP-T1 | Aci6 | - |
| LT984690.1 | SAMEA104446236 | K50 | p2K50 | 79598 | Rep | RP-T1 | Aci6 | SPC58401.1 |
| MG954377.1 | SAMN14228297 | SGH9601 | pS21-2 | 123432 | Rep | RP-T1 | - | AWO68424.1 |
| MG954379.1 | SAMN14228295 | SGH0905 | pS32-2 | 70833 | Rep | RP-T1 | Aci6 | AWO68573.1 |
| MK243454.1 | SAMN14227902 | 09A16CRGN0014 | pCRA914-67 | 66886 | Rep | RP-T1 | Aci6 | QAR16211.1 |
| MK386681.1 | SAMN14228691 | ABAY09008 | pABAY09008_1B | 74241 | Rep | RP-T1 | Aci6 | QBN23166.1 |
| MK531538.1 | - | MC23 | pMC23.1 | 67441 | Rep | RP-T1 | - | QCO89676.1 |
| CP003501.1 | SAMN02603104 | MDR-TJ | pABTJ1 | 77528 | Rep | RP-T2 | - | - |
| CP003887.1 | SAMN02604244 | BJAB07104 | p1BJAB07104 | 70170 | Rep | RP-T2 | - | - |
| CP003888.1 | SAMN02604246 | BJAB0868 | p2BJAB0868 | 70167 | Rep | RP-T2 | - | - |
| CP018144.1 | SAMN06046790 | HRAB-85 | pHRAB-85 | 77513 | Rep | RP-T2 | - | APF45706.1 |
| CP018422.1 | SAMN06109232 | XDR-BJ83 | pBJ83 | 69069 | Rep | RP-T2 | - | APM50974.1 |
| KM922672.1 | SAMN03103694 | A221 | pAZJ221 | 77530 | Rep | RP-T2 | - | - |
| MK386682.1 | SAMN14228690 | ABAY10001 | pABAY10001_1C | 54627 | Rep | RP-T2 | - | QBN23313.1 |
| CP013925.1 | SAMN03941550 | KBN10P02143 | pKBN10P02143 | 52517 | Rep | RP-T3 | - | ALY01450.1 |
| CM009039.2 | SAMN08093365 | ZQ6 | p2ZQ6 | 6772 | Rep | RP-T4 | - | PQL72098.1 |
| CP042563.1 | SAMN12289292 | E47 | pE47_007 | 4715 | Rep | RP-T5 | - | QFH47718.1 |
| CP059390.1 | SAMN15541804 | 36-1512 | p4.36-1512 | 4721 | Rep | RP-T5 | - | QLY88352.1 |

**Table S3** Rep_3 plasmids

| Acession number | Biosample | strain_name | Plasmid name | Length(bp) | Rep_family | Type | Bertini scheme | Locus/Protein_id |
|---|---|---|---|---|---|---|---|---|
| CP000523.1 | SAMN02604331 | ATCC 17978 | pAB2 | 11302 | Rep_3 | R3-T1 | A1S_3472 | ABO13861.1 |
| CU459138.1 | SAMEA3138279 | AYE | p2ABAYE | 9661 | Rep_3 | R3-T1 | p2ABAYE0002 | CAM84615.1 |
| CP000523.1 | SAMN02604331 | ATCC 17978 | pAB2 | 11302 | Rep_3 | R3-T1 | Aci1 | ABO13861.1 |
| CP001183.2 | SAMN02603051 | AB0057 | pAB0057 | 8731 | Rep_3 | R3-T1 | Aci1 | ACJ43223.1 |
| CP001183.2 | SAMN02603051 | AB0057 | pAB0057 | 8731 | Rep_3 | R3-T1 | Aci1 | ACJ43223.1 |
| CP024419.1 | SAMN07736509 | A388 | pA388 | 33036 | Rep_3 | R3-T1 | Aci1 | ATP89005.1 |
| CP031381.2 | SAMN09302593 | ACICU | pACICU1b | 24268 | Rep_3 | R3-T1 | Aci1 | QCS03991.1 |
| CU459138.1 | SAMEA3138279 | AYE | p2ABAYE | 9661 | Rep_3 | R3-T1 | Aci1 | CAM84615.1 |
| MN266872.1 | - | N/A | pAC1-BRL | 16673 | Rep_3 | R3-T1 | Aci1 | QHW11277.1 |
| AB823544.1 | SAMN14229301 | NCGM 253 | pAB-NCGM253 | 8970 | Rep_3 | R3-T1 | Aci1 | - |
| AEOY01000096.1 | SAMN02471598 | 3990 | p2ABST2 | 21846 | Rep_3 | R3-T1 | Aci1 | - |
| AEOZ01000236.1 | SAMN02471606 | 3909 | p1ABST78 | 26411 | Rep_3 | R3-T1 | Aci1 | - |
| AEPA01000396.1 | SAMN02471587 | 4190 | p2ABST25 | 8970 | Rep_3 | R3-T1 | Aci1 | - |
| AFDN01000003.1 | SAMN02436468 | Canada BC-5 | pCanadaBC5-8.7 | 8731 | Rep_3 | R3-T1 | Aci1 | EJO35917.1 |
| AYEX01000118.1 | SAMN06650245 | CBA7 | pABUH6a-8.8 | 8763 | Rep_3 | R3-T1 | Aci1 | ETR37079.1 |
| CM003314.1 | SAMN02906929 | MRSN 7339 | pMRSN7339-8.7 | 8731 | Rep_3 | R3-T1 | Aci1 | KLT75075.1 |
| CM003317.1 | SAMN02906928 | MRSN 58 | pMRSN58-8.7 | 8731 | Rep_3 | R3-T1 | Aci1 | KLT95674.1 |
| CM003741.1 | SAMN04407353 | MEX11594 | p2MEX11594 | 5557 | Rep_3 | R3-T1 | Aci1 | - |
| CM003909.1 | SAMN03450127 | AB210M | pAB0057 | 8781 | Rep_3 | R3-T1 | Aci1 | KZC87498.1 |
| CM009038.2 | SAMN08093365 | ZQ6 | p1ZQ6 | 8905 | Rep_3 | R3-T1 | Aci1 | PQL72050.1 |
| CM009043.2 | SAMN08093364 | ZQ5 | p2ZQ5 | 8731 | Rep_3 | R3-T1 | Aci1 | PQL83867.1 |
| CM009924.1 | SAMN07602915 | CCUG 70743 | pAba70743_1 | 10880 | Rep_3 | R3-T1 | Aci1 | PXF35729.1 |
| CP002523.1 | SAMN02603891 | TCDC-AB0715 | p1ABTCDC0715 | 8731 | Rep_3 | R3-T1 | Aci1 | ADX94286.1 |
| CP003850.1 | SAMN02604246 | BJAB0868 | p1BJAB0868 | 8721 | Rep_3 | R3-T1 | Aci1 | AGQ12262.1 |
| CP006964.1 | SAMN03081512 | AB07 | pPKAB07 | 8805 | Rep_3 | R3-T1 | Aci1 | AHJ95281.1 |
| CP007550.1 | SAMN02471420 | AC12 | pAC12 | 8731 | Rep_3 | R3-T1 | Aci1 | AHX30527.1 |
| CP007578.1 | SAMN02471421 | AC30 | pAC30a | 8685 | Rep_3 | R3-T1 | Aci1 | AHX67213.1 |
| CP008708.1 | SAMN02894434 | AB5075-UW | p2AB5075 | 8731 | Rep_3 | R3-T1 | Aci1 | AKA33680.1 |
| CP008850.1 | SAMN02714232 | AC29 | pAC29a | 8737 | Rep_3 | R3-T1 | Aci1 | AKB09309.1 |
| CP010782.1 | SAMN03248539 | A1 | pA1-1 | 8731 | Rep_3 | R3-T1 | Aci1 | AJF83584.1 |
| CP012955.1 | SAMN04029125 | D36 | pD36-3 | 9276 | Rep_3 | R3-T1 | Aci1 | ALJ89812.1 |
| CP014217.1 | SAMN04448797 | YU-R612 | p1YU-R612 | 5465 | Rep_3 | R3-T1 | Aci1 | AMC17828.1 |
| CP014293.1 | SAMN04457967 | AB34299 | p1AB34299 | 15645 | Rep_3 | R3-T1 | Aci1 | AQU58924.1 |
| CP015486.1 | SAMN03277095 | ORAB01 | pORAB01-3 | 15198 | Rep_3 | R3-T1 | - | ANB90648.1 |
| CP020576.1 | SAMN06650263 | SSA12 | pSSA12_2 | 8730 | Rep_3 | R3-T1 | Aci1 | ARF94646.1 |
| CP021786.1 | SAMN07125723 | A85 | pA85-2 | 8731 | Rep_3 | R3-T1 | Aci1 | AHM95263.1 |
| CP026708.1 | SAMN04014897 | AR_0056 | tig00000534_pilon | 8731 | Rep_3 | R3-T1 | Aci1 | AVE48090.1 |
| CP027124.1 | SAMN04014897 | AR_0056 | p1AR_0056 | 8731 | Rep_3 | R3-T1 | Aci1 | AVN07781.1 |
| CP027244.1 | SAMN08364584 | WCHAB005078 | p2_005078 | 8731 | Rep_3 | R3-T1 | Aci1 | AVN12717.1 |
| CP027529.1 | SAMN04014924 | AR_0083 | pAR_0083 | 8731 | Rep_3 | R3-T1 | Aci1 | AVN27928.1 |
| CP027609.1 | SAMN04014943 | AR_0102 | p1AR_0102 | 8731 | Rep_3 | R3-T1 | - | - |
| CP031446.1 | SAMN09769497 | MDR-UNC | pAB120 | 10879 | Rep_3 | R3-T1 | Aci1 | QBA07847.1 |
| CP033870.1 | SAMN10411605 | MRSN15313 | p597A-6.7, | 6667 | Rep_3 | R3-T1 | Aci1 | AYY91181.1 |
| CP040261.1 | SAMN11621520 | P7774 | p2P7774 | 14880 | Rep_3 | R3-T1 | Aci1 | QCR91174.1 |
| CP050387.1 | SAMN14409516 | VB82 | pVB82_2 | 7540 | Rep_3 | R3-T1 | Aci1 | QJH23945.1 |
| CP050391.1 | SAMN14409813 | VB723 | pVB723_1 | 8731 | Rep_3 | R3-T1 | Aci1 | QJH16202.1 |
| CP050393.1 | SAMN14410107 | VB11737 | pVB11737_2 | 8731 | Rep_3 | R3-T1 | Aci1 | QJH08912.1 |
| CP050402.1 | SAMN14410138 | VB2181 | pVB2181 | 8731 | Rep_3 | R3-T1 | Aci1 | QJH08812.1 |
| CP050411.1 | SAMN14420246 | PM1912235 | pPM192235_1 | 8731 | Rep_3 | R3-T1 | Aci1 | QJG93743.1 |
| CP050414.1 | SAMN14420253 | PM192696 | pPM192696_2 | 8732 | Rep_3 | R3-T1 | Aci1 | QJG90135.1 |
| CP050419.1 | SAMN14420254 | PM193665 | pPM193665_4 | 7540 | Rep_3 | R3-T1 | Aci1 | QJG86381.1 |
| CP050422.1 | SAMN14415334 | VB2200 | pVB2200_1 | 8731 | Rep_3 | R3-T1 | Aci1 | QJG97436.1 |
| CP050429.1 | SAMN14420255 | PM194188 | pPM194122_4 | 7549 | Rep_3 | R3-T1 | Aci1 | QJG82479.1 |
| CP050524.1 | SAMN14410111 | VB7036 | pVB7036_1 | 8730 | Rep_3 | R3-T1 | Aci1 | QJG74742.1 |
| CP050527.1 | SAMN14414747 | VB2139 | pVB2139_1 | 8731 | Rep_3 | R3-T1 | Aci1 | QJG70972.1 |
| CP050906.1 | SAMN14308892 | DT-Ab057 | p1DT-Ab057 | 8731 | Rep_3 | R3-T1 | Aci1 | QIX32739.1 |
| CP050910.1 | SAMN14308866 | DT-Ab022 | p1DT-Ab0 | 8731 | Rep_3 | R3-T1 | Aci1 | QIX36700.1 |
| CP050913.1 | SAMN14308864 | DT-Ab020 | p1DT-Ab020 | 8731 | Rep_3 | R3-T1 | Aci1 | QIX40567.1 |
| CP050918.1 | SAMN14308849 | DT-Ab003 | p1DT-Ab003 | 8731 | Rep_3 | R3-T1 | Aci1 | QIX48098.1 |
| CP051475.1 | SAMN14414778 | VB2107 | pVB2107_1 | 8731 | Rep_3 | R3-T1 | Aci1 | QJH01183.1 |
| CP056785.1 | SAMN15344688 | TP1 | pTP1A | 8731 | Rep_3 | R3-T1 | Aci1 | QLA74147.1 |
| CP058626.1 | SAMN15437745 | ATCC BAA1605 | pATCCBAA1605 | 8731 | Rep_3 | R3-T1 | Aci1 | QLG82497.1 |
| CP060012.1 | SAMN15735522 | TP2 | pTP2A | 8731 | Rep_3 | R3-T1 | Aci1 | QPO01560.1 |
| CP060014.1 | SAMN15738014 | TP3 | pTP3A | 8731 | Rep_3 | R3-T1 | Aci1 | QPO05067.1 |
| CP061526.1 | SAMN12391537 | CFSAN093705 | pCFSAN093705 | 14639 | Rep_3 | R3-T1 | - | QNV23750.1 |
| CP066017.1 | SAMN16357205 | FDAARGOS_1036 | pFDAARGOS_1036 | 10308 | Rep_3 | R3-T1 | Aci1 | QQB69061.1 |
| CP066231.1 | SAMN17073618 | G20AB011 | pG20AB011-2, | 8731 | Rep_3 | R3-T1 | Aci1 | QQD98226.1 |
| CP066240.1 | SAMN17073615 | G20AB007 | pG20AB007-3 | 8731 | Rep_3 | R3-T1 | Aci1 | QQE02022.1 |
| CP069842.1 | SAMN16357502 | FDAARGOS_1360 | p1FDAARGOS_1360 | 14879 | Rep_3 | R3-T1 | Aci1 | QRR53505.1 |
| CP069852.1 | SAMN16357501 | FDAARGOS_1359 | pFDAARGOS_1359 | 8731 | Rep_3 | R3-T1 | Aci1 | QRR71166.1 |
| CP071920.1 | SAMN18276099 | GIMC5510 | pABT-897-17 | 13480 | Rep_3 | R3-T1 | Aci1 | QTF98049.1 |
| CP072125.1 | SAMN18396008 | KSK1 | p3KSK1 | 7540 | Rep_3 | R3-T1 | Aci1 | QTH58467.1 |
| CP072273.1 | SAMN18452234 | KSK6 | p3KSK6 | 7540 | Rep_3 | R3-T1 | Aci1 | QTK45779.1 |
| CP072278.1 | SAMN18452334 | KSK7 | p3KSK7 | 7540 | Rep_3 | R3-T1 | Aci1 | QTK62150.1 |
| CP072283.1 | SAMN18452342 | KSK10 | p3KSK10 | 7540 | Rep_3 | R3-T1 | Aci1 | QTK54004.1 |
| CP072288.1 | SAMN18452659 | KSK11 | p3KSK11 | 7540 | Rep_3 | R3-T1 | Aci1 | QTK58080.1 |
| CP072293.1 | SAMN18452698 | KSK18 | p3KSK18 | 7540 | Rep_3 | R3-T1 | Aci1 | QTK66220.1 |
| CP072298.1 | SAMN18452699 | KSK19 | p3KSK19 | 7540 | Rep_3 | R3-T1 | Aci1 | QTK70313.1 |
| CP072303.1 | SAMN18452712 | KSK20 | p3KSK20 | 7540 | Rep_3 | R3-T1 | Aci1 | QTK74379.1 |
| CP072307.1 | SAMN18452713 | KSK Sensitive | p2KSKSensitive | 6520 | Rep_3 | R3-T1 | - | QTK77975.1 |
| CP072401.1 | SAMN18451305 | KSK2 | p3KSK2 | 7540 | Rep_3 | R3-T1 | Aci1 | QTL10389.1 |
| HG380023.1 | SAMN14229284 | 107m | ABIBUN107mP1 | 8731 | Rep_3 | R3-T1 | Aci1 | CDG34533.1 |
| JACGEJ010000130.1 | SAMN15501050 | AbCTX19 | pAbCTX19_9kb | 8970 | Rep_3 | R3-T1 | Aci1 | MBL4063611.1 |
| JHUI01000005.1 | SAMN06650264 | 15A5 | pAB5075 | 8819 | Rep_3 | R3-T1 | Aci1 | KGP67530.1 |
| JX069966.1 | SAMN14226317 | K60 | pAB120 | 10879 | Rep_3 | R3-T1 | Aci1 | AFO83979.1 |
| KJ586856.1 | SAMN14226822 | G7 | pAB-G7-1, | 8731 | Rep_3 | R3-T1 | Aci1 | AHM95359.1 |
| KR535993.1 | SAMN14226627 | A105 | pA105-2 | 9830 | Rep_3 | R3-T1 | Aci1 | ALN43409.1 |
| KU869528.1 | SAMN14227234 | A297(RUH875) | pA297-2 | 8731 | Rep_3 | R3-T1 | Aci1 | AMX23364.1 |
| KX230793.1 | SAMN14227146 | MAL | pMAL-1 | 9810 | Rep_3 | R3-T1 | Aci1 | ANR95803.1 |
| KY202456.1 | SAMN14227361 | AB1433 | pIBAC_oxa58_1433 | 26496 | Rep_3 | R3-T1 | Aci1 | ARD69932.1 |
| KY202457.1 | SAMN14227360 | AB2RED09 | pIBAC_oxa58_2RED | 25311 | Rep_3 | R3-T1 | Aci1 | ARD69954.1 |
| KY202458.1 | SAMN14227359 | AB20C15 | pIBAC_oxa58_20C15 | 26781 | Rep_3 | R3-T1 | Aci1 | ARD69975.1 |
| MG954378.1 | SAMN14228296 | SGH0905 | pS32-1 | 13545 | Rep_3 | R3-T1 | Aci1 | AWO68558.1 |
| MH362811.1 | SAMN07258672 | 11A1314CRGN088 | pO237-3 | 18475 | Rep_3 | R3-T1 | - | AXF37697.1 |
| MH362812.1 | SAMN07258655 | 11A1213CRGN008 | pO237-1 | 15199 | Rep_3 | R3-T1 | - | AXF37710.1 |
| MH362813.1 | SAMN07258647 | 11A1213CRGN055 | pO237-2 | 18475 | Rep_3 | R3-T1 | - | AXF37717.1 |
| MK386683.1 | SAMN14228689 | ABAY14012 | pABAY14012_4D | 8753 | Rep_3 | R3-T1 | Aci1 | QBN23334.1 |
| MK431775.1 | SAMN14229072 | 11A14CRGN003 | pO237-4 | 15199 | Rep_3 | R3-T1 | - | QBQ02584.1 |
| MK531537.1 | - | MC1/MC23 | pMC1.2/pMC23.2 | 8731 | Rep_3 | R3-T1 | Aci1 | QCW06040.1 |
| CU468232.1 | SAMEA3138277 | SDF | p2ABSDF | 25104 | Rep_3 | R3-T10 | p2ABSDF0001 | CAP02944.1 |

| | | | | | | | | |
|---|---|---|---|---|---|---|---|---|
| CU468232.1 | SAMEA3138277 | SDF | p2ABSDF | 25014 | Rep_3 | R3-T10 | p2ABSDF0001 | CAP02944.1 |
| LR026972.1 | SAMEA4646212 | RDK36_28 | pKCRI-28-1 | 29606 | Rep_3 | R3-T10 | p2ABSDF0001 | - |
| CM003742.1 | SAMN04407353 | MEX11594 | p1MEX11594 | 4437 | Rep_3 | R3-T10 | p2ABSDF0001 | - |
| CP051871.1 | SAMN14667516 | Ab-D10a-a | pAb-D10a-a_2 | 8495 | Rep_3 | R3-T10 | p2ABSDF0001 | QJF33703.1 |
| CP051877.1 | SAMN14667515 | Ab-B004d-c | pAb-B004d-c_2 | 8495 | Rep_3 | R3-T10 | p2ABSDF0001 | QJF37590.1 |
| CP059730.1 | SAMN15501316 | AbCTX13 | pAbCTX13_7kb | 7055 | Rep_3 | R3-T10 | p2ABSDF0001 | QRN23772.1 |
| JACGEK010000079.1 | SAMN15501052 | AbCTX17 | pAbCTX17_7kb | 7055 | Rep_3 | R3-T10 | p2ABSDF0001 | MBL4076741.1 |
| LT984691.1 | SAMEA104446236 | K50 | p1K50 | 9539 | Rep_3 | R3-T10 | p2ABSDF0001 | SPC58512.1 |
| GU978998.1 | - | - | p844 | 1119 | Rep_3 | R3-T11 | Aci4 | ADM89093.1 |
| CM009651.1 | SAMN08093362 | ZQ3 | p5ZQ3 | 16470 | Rep_3 | R3-T11 | Aci4 | PQJ03708.1 |
| CP040045.1 | SAMN11554497 | VB958 | p1VB958 | 16485 | Rep_3 | R3-T11 | Aci4 | QCP18714.1 |
| CP040049.1 | SAMN11554995 | VB1190 | pVB1190 | 16479 | Rep_3 | R3-T11 | Aci4 | QCP22235.1 |
| CP040055.1 | SAMN11557490 | VB35179 | p1VB35179 | 16467 | Rep_3 | R3-T11 | Aci4 | QCP25971.1 |
| CP050389.1 | SAMN14409628 | VB473 | pVB473_1 | 16470 | Rep_3 | R3-T11 | Aci4 | QJH19807.1 |
| CP050405.1 | SAMN14414761 | VB2486 | pVB2486_2 | 14906 | Rep_3 | R3-T11 | Aci4 | QJH05136.1 |
| CP050418.1 | SAMN14420254 | PM193665 | pPM193665_3 | 18783 | Rep_3 | R3-T11 | Aci4 | QJG86361.1 |
| CP050427.1 | SAMN14420255 | PM194188 | pPM194122_2 | 18783 | Rep_3 | R3-T11 | Aci4 | QJG82448.1 |
| AFDB02000003.1 | SAMN00114928 | Naval-81 | pNaval81-26 | 26089 | Rep_3 | R3-T12 | - | EJP56825.1 |
| CM009030.2 | SAMN08093369 | ZQ10 | p1ZQ10 | 35194 | Rep_3 | R3-T12 | - | PQJ03477.1 |
| CM009083.3 | SAMN08093368 | ZQ9 | p1ZQ9 | 35194 | Rep_3 | R3-T12 | - | PQJ03454.1 |
| AFDA02000011.1 | SAMN00114927 | Naval-18 | pNaval18-7.0 | 7032 | Rep_3 | R3-T12 | - | EJP48327.1 |
| ALII01000018.1 | SAMN02436485 | IS-123 | pIS123-12 | 11600 | Rep_3 | R3-T12 | - | EJO37651.1 |
| CP045108.1 | SAMN12389466 | ATCC 19606 | p1ATCC19606 | 7655 | Rep_3 | R3-T12 | - | QFQ03441.1 |
| CP065888.1 | SAMN13450447 | FDAARGOS_917 | p1FDAARGOS_917 | 6778 | Rep_3 | R3-T12 | - | QQA28272.1 |
| AFDA02000008.1 | SAMN00114927 | Naval-18 | pNaval18-8.4 | 8422 | Rep_3 | R3-T13 | - | EJP48482.1 |
| CP010400.1 | SAMN03263969 | 6200 | p6200-9.327kb | 9327 | Rep_3 | R3-T13 | - | AJB69037.1 |
| CP026126.1 | SAMN06040401 | ABNIH28 | pABA-6973 | 11305 | Rep_3 | R3-T13 | - | AUT39979.1 |
| CP044521.1 | SAMN12859885 | 29FS20 | p29FS20-2 | 12731 | Rep_3 | R3-T13 | - | QLF12502.1 |
| CP051865.1 | SAMN14667518 | Ab-C102 | pAb-C102_3 | 19853 | Rep_3 | R3-T13 | - | QJF29816.1 |
| CP059388.1 | SAMN15541804 | 36-1512 | p2.36-1512 | 9458 | Rep_3 | R3-T13 | - | QLY88340.1 |
| CP033871.1 | SAMN10411605 | MRSN15313 | p597A-14.8 | 14087 | Rep_3 | R3-T14 | - | AYY91191.1 |
| MG100202.1 | SAMN14228542 | Ab825 | pAb825_36 | 35743 | Rep_3 | R3-T14 | - | AVR61203.1 |
| MN266872.1 | - | N/A | pAC1-BRL | 16673 | Rep_3 | R3-T14 | - | QHW11287.1 |
| KY984045.1 | SAMN07509424 | AB242 | pAb242_9 | 9284 | Rep_3 | R3-T14 | - | AUO31880.1 |
| MG520098.1 | SAMN14228448 | Ab244 | pAb244_7 | 7965 | Rep_3 | R3-T14 | - | AVX50861.1 |
| MK323042.1 | SAMN14228622 | Acb-45063 | pAb45063_a | 19808 | Rep_3 | R3-T14 | - | QBK17989.1 |
| MK531541.1 | - | MC75 | pMC75.2 | 13903 | Rep_3 | R3-T14 | - | QCO89772.1 |
| CP062923.1 | SAMN16304032 | Res13-Abat-PEA21-P4-01-A | p1Res13-Abat | 5242 | Rep_3 | R3-T15 | - | QPF15413.1 |
| AFDL01000005.1 | SAMN02436631 | OIFC143 | pOIFC143-2.3 | 2277 | Rep_3 | R3-T15 | - | EJG16531.1 |
| CM009646.1 | SAMN08093369 | ZQ10 | p3ZQ10 | 2277 | Rep_3 | R3-T15 | - | PQJ03470.1 |
| JADAIY010000264.1 | SAMN16304042 | Res13-Abat-PEA28-P5-02-A | pRes13-Abat-PEA28-P5-02-A | 2279 | Rep_3 | R3-T15 | - | - |
| MK386684.1 | SAMN14228688 | ABAY15001 | pABAY15001_6E | 2278 | Rep_3 | R3-T15 | - | QBN23345.1 |
| CM016516.1 | SAMN10261616 | TG31307 | pTG31307 | 55269 | Rep_3 | R3-T16 | - | THJ58480.1 |
| CP039342.1 | SAMN10261613 | TG31986 | pTG31986 | 55269 | Rep_3 | R3-T16 | - | QCD20890.1 |
| CP039344.1 | SAMN10261535 | TG31302 | pTG31302 | 55268 | Rep_3 | R3-T16 | - | QCD24650.1 |
| CP039932.1 | SAMN10261537 | TG29392 | pTG29392_2 | 55269 | Rep_3 | R3-T16 | - | QCO80749.1 |
| CP072306.1 | SAMN18452713 | KSK Sensitive | p1KSKSensitive | 55356 | Rep_3 | R3-T16 | - | QTK77957.1 |
| CP015365.1 | SAMN04485290 | 3207 | pAba3207a | 13478 | Rep_3 | R3-T17 | - | ANC38759.1 |
| CP022284.1 | SAMN07289440 | 7804 | pAba7804a | 12381 | Rep_3 | R3-T17 | - | ASO73059.1 |
| CP023032.1 | SAMN07284119 | 7847 | pAba7847a | 13478 | Rep_3 | R3-T17 | - | AXW92573.1 |
| CP033244.1 | SAMN07520235 | 7835 | pAba7835a | 8536 | Rep_3 | R3-T17 | - | QFY70909.1 |
| CP050428.1 | SAMN14420255 | PM194188 | pPM194122_3 | 7695 | Rep_3 | R3-T17 | - | QJG82477.1 |
| CU459137.1 | SAMEA3138279 | AYE | p1ABAYE | 5644 | Rep_3 | R3-T18 | p1ABAYE0001 | CAM84608.1 |
| CU459137.1 | SAMEA3138279 | AYE | p1ABAYE | 5644 | Rep_3 | R3-T18 | p1ABAYE0001 | CAM84608.1 |
| CM004453.1 | SAMN03699770 | M3AC14-8 | p1M3AC14-8 | 5441 | Rep_3 | R3-T18 | p1ABAYE0001 | OBR16604.1 |
| CP042561.1 | SAMN12289292 | E47 | pE47_005 | 5234 | Rep_3 | R3-T18 | p1ABAYE0001 | QFH47707.1 |
| CP046901.1 | SAMN13565236 | A1429 | pA1429a | 7852 | Rep_3 | R3-T18 | p1ABAYE0001 | QLB37634.1 |
| CP024419.1 | SAMN07736509 | A388 | pA388 | 33036 | Rep_3 | R3-T19 | Aci10 | ATP89028.1 |
| CP031381.2 | SAMN09302593 | ACICU | pACICU1b | 24268 | Rep_3 | R3-T19 | Aci10 | QCS03997.1 |
| CP027179.1 | SAMN04014911 | AR_0070 | p2AR_0070 | 39535 | Rep_3 | R3-T19 | Aci10 | AVI35083.1 |
| CP027184.1 | SAMN04014893 | AR_0052 | p3AR_0052 | 60699 | Rep_3 | R3-T19 | Aci10 | AVI39286.1 |
| FM210331.1 | SAMN14229501 | VA-566/00 | pABVA01 | 8963 | Rep_3 | R3-T2 | Aci2 | CAR65318.1 |
| FM210331.1 | SAMN14229501 | VA-566/00 | pABVA01 | 8963 | Rep_3 | R3-T2 | Aci2 | CAR65318.1 |
| MN495625.1 | - | A2485 | pA2485 | 15405 | Rep_3 | R3-T2 | Aci2 | QID24189.1 |
| MN495626.1 | - | A2503 | pA2503 | 15405 | Rep_3 | R3-T2 | Aci2 | QID24208.1 |
| AP022240.1 | SAMD00194616 | WP8-W18-ESBL-11 | pWP8-W18-ESBL-11_2 | 10735 | Rep_3 | R3-T2 | - | - |
| AYFI01000019.1 | SAMN07736509 | A388 | pABUH6b-10 | 10030 | Rep_3 | R3-T2 | Aci2 | ETR11343.1 |
| CM009045.2 | SAMN08093363 | ZQ4 | p1ZQ4 | 8331 | Rep_3 | R3-T2 | Aci2 | PST50036.1 |
| CP026748.1 | SAMN08364585 | WCHAB005133 | p1_005133 | 5602 | Rep_3 | R3-T2 | Aci2 | AVE88593.1 |
| CP029573.1 | SAMN06650239 | HWBA8 | pDA33098-9 | 8963 | Rep_3 | R3-T2 | Aci2 | AWO18631.1 |
| CP035044.1 | SAMN05238571 | ABUH796 | p13.0Kbp | 12952 | Rep_3 | R3-T2 | Aci2 | QAS95996.1 |
| CP035048.1 | SAMN05238672 | ABUH793 | p10.9Kbp | 10945 | Rep_3 | R3-T2 | Aci2 | QAS99819.1 |
| CP035050.1 | SAMN05238697 | ABUH773 | p11.8Kbp | 11810 | Rep_3 | R3-T2 | Aci2 | QAT03382.1 |
| CP035053.1 | SAMN05238628 | YU-R612 | p11.0Kbp | 10967 | Rep_3 | R3-T2 | Aci2 | QAT07114.1 |
| CP039521.1 | SAMN10261584 | TG22627 | pTG22627 | 5602 | Rep_3 | R3-T2 | Aci2 | QCH38679.1 |
| CP039995.1 | SAMN10261544 | TG22182 | pTG22182_2 | 5602 | Rep_3 | R3-T2 | Aci2 | QCO84567.1 |
| CP040427.1 | SAMN11660471 | PB364 | pPB364_2 | 10967 | Rep_3 | R3-T2 | Aci2 | QCT18126.1 |
| CP041589.1 | SAMN12158047 | J9 | pJ9-2 | 10967 | Rep_3 | R3-T2 | Aci2 | QDM68406.1 |
| CP042208.1 | SAMN08637743 | DS002 | pTS11291 | 11291 | Rep_3 | R3-T2 | Aci2 | QDX16375.1 |
| CP042560.1 | SAMN12289292 | E47 | pE47_004 | 7703 | Rep_3 | R3-T2 | Aci2 | QFH47694.1 |
| CP044358.1 | SAMN12825295 | CAM180-1 | pCAM180B | 16096 | Rep_3 | R3-T2 | Aci2 | QEY06144.1 |
| CP047976.1 | SAMN13884837 | DETAB-P2 | pDETAB3 | 9132 | Rep_3 | R3-T2 | Aci2 | QMS84177.1 |
| CP050434.1 | SAMN14422682 | PM194229 | pPM194229_2 | 10697 | Rep_3 | R3-T2 | Aci2 | QJG78557.1 |
| CP059389.1 | SAMN15541804 | 36-1512 | p3.36-1512 | 5688 | Rep_3 | R3-T2 | Aci2 | QLY88346.1 |
| CP067103.1 | SAMN12399660 | ATCC BAA-1790 | pNC2 | 10955 | Rep_3 | R3-T2 | Aci2 | QQY87247.1 |
| GQ904227.1 | SAMN14225470 | - | pMMCU3 | 8964 | Rep_3 | R3-T2 | Aci2 | ADB23472.1 |
| HG977525.1 | SAMEA3158456 | CS01 | pCS01C | 8174 | Rep_3 | R3-T2 | Aci2 | - |
| HG977529.1 | SAMEA3158506 | CR17 | pCR17C | 8047 | Rep_3 | R3-T2 | Aci2 | - |
| KJ534568.1 | SAMN14226773 | ATCC 223 | AbATCC223 | 8840 | Rep_3 | R3-T2 | Aci2 | AIA61624.1 |
| KJ534569.1 | SAMN14226772 | ATCC 329 | AbATCC329 | 8842 | Rep_3 | R3-T2 | Aci2 | AIA61634.1 |
| KM051986.1 | SAMN14226464 | D72 | pD72-1 | 10967 | Rep_3 | R3-T2 | Aci2 | AIH07953.1 |
| MG954376.1 | SAMN14228298 | SGH9601 | pS21-1 | 12952 | Rep_3 | R3-T2 | Aci2 | AWO68412.1 |
| CP051870.1 | SAMN14667516 | Ab-D10a-a | pAb-D10a-a_1 | 48239 | Rep_3 | R3-T20 | - | QJF33663.1 |
| CP051876.1 | SAMN14667515 | Ab-B004d-c | pAb-B004d-c_1 | 48239 | Rep_3 | R3-T20 | - | QJF37552.1 |
| CP053220.1 | SAMN14833556 | DT01139C | p2DT01139C | 63650 | Rep_3 | R3-T20 | - | QLI41749.1 |
| KY216144.1 | - | RCH51 | pRCH51-3 | 52789 | Rep_3 | R3-T20 | - | AQT19035.1 |
| CP026129.1 | SAMN06040401 | ABNIH28 | pABA-2f10 | 130044 | Rep_3 | R3-T21 | - | AUT40247.1 |
| CP026749.2 | SAMN08364585 | WCHAB005133 | pOXA58_005133, | 42455 | Rep_3 | R3-T21 | - | AVE88635.1 |
| CP038501.1 | SAMN11298775 | CIAT758 | p3CIAT758 | 78125 | Rep_3 | R3-T21 | - | QBY16306.1 |
| CP053219.1 | SAMN14833556 | DT01139C | p3DT01139C | 97161 | Rep_3 | R3-T21 | - | QLI41705.1 |
| CM013137.1 | SAMN10662617 | AB18PR065 | pAB18PR065-MCR-4.3 | 25602 | Rep_3 | R3-T22 | - | RUT37677.1 |
| CP033872.1 | SAMN10411605 | MRSN15313 | pAb-MCR4.1 | 35502 | Rep_3 | R3-T22 | - | AYY91210.1 |

| | | | | | | | | |
|---|---|---|---|---|---|---|---|---|
| CP038261.1 | SAMN10386508 | 39741 | pEH_mcr4.3 | 18786 | Rep_3 | R3-T22 | - | QBR82861.1 |
| CP038265.1 | SAMN10386510 | LEV1449/17Ec | pEC_mcr4.3 | 43093 | Rep_3 | R3-T22 | - | QBR79279.1 |
| AFDB02000005.1 | SAMN00114928 | Naval-81 | pNaval81-13 | 12634 | Rep_3 | R3-T23 | - | - |
| AFDO01000021.1 | SAMN02436551 | Naval-17 | pNaval17-13 | 12636 | Rep_3 | R3-T23 | - | EJG28245.1 |
| CM009033.2 | SAMN08093367 | ZQ8 | p1ZQ8 | 12636 | Rep_3 | R3-T23 | - | PQL85636.1 |
| CP050406.1 | SAMN14414761 | VB2486 | pVB2486_3 | 12574 | Rep_3 | R3-T23 | - | QJH05158.1 |
| CP033871.1 | SAMN10411605 | MRSN15313 | p597A-14.8 | 14087 | Rep_3 | R3-T24 | - | AYY91202.1 |
| CP034097.1 | SAMN10441121 | A52 | pA52-OXA-72 | 8493 | Rep_3 | R3-T24 | - | QAB42528.1 |
| CP043954.1 | SAMN12769618 | K09-14 | pK09-14 | 7791 | Rep_3 | R3-T24 | - | QER77251.1 |
| KY704308.1 | SAMN14227859 | IHIT32296 | pAbIHIT32296 | 8493 | Rep_3 | R3-T24 | - | ASN73624.1 |
| CP012956.1 | SAMN04029125 | D36 | pD36-4 | 47457 | Rep_3 | R3-T25 | - | ALJ89824.1 |
| CP051863.1 | SAMN14667518 | Ab-C102 | pAb-C102_1 | 90089 | Rep_3 | R3-T25 | - | QJF29769.1 |
| CP051867.1 | SAMN14667517 | Ab-C63 | pAb-C63_1 | 81353 | Rep_3 | R3-T25 | - | QJF41224.1 |
| AYFH01000057.1 | SAMN02597386 | UH7607 | pABUH2b-5.4 | 5355 | Rep_3 | R3-T26 | - | ETR11099.1 |
| CP026340.1 | SAMN07559626 | 810CP | pAba810CPa | 5281 | Rep_3 | R3-T26 | - | AXG87071.1 |
| CP044518.1 | SAMN12860376 | 31FS3-2 | p31FS3-2-1 | 6099 | Rep_3 | R3-T26 | - | QLF08691.1 |
| CP034094.1 | SAMN10441121 | A52 | pA52-2 | 27452 | Rep_3 | R3-T27 | - | QAB42494.1 |
| CP051864.1 | SAMN14667518 | Ab-C102 | pAb-C102_2 | 67097 | Rep_3 | R3-T27 | - | QJF29792.1 |
| CP059478.1 | SAMN15637465 | 17-84 | p17-84_OXA | 108715 | Rep_3 | R3-T28 | - | QNB01597.1 |
| CP047975.1 | SAMN13884837 | DETAB-P2 | pDETAB2 | 100072 | Rep_3 | R3-T28 | - | QMS84089.1 |
| LN833432.1 | SAMEA3298506 | CHI-32 | pNDM-32 | 84623 | Rep_3 | R3-T28 | - | - |
| CP062922.1 | SAMN16304032 | Res13-Abat-PEA21-P4-01-A | p2Res13-Abat | 14288 | Rep_3 | R3-T29 | - | QPF15404.1 |
| JADAJJ010000299.1 | SAMN16304029 | Res13-Abat-PEA16-P5-01-A | pRes13-Abat-PEA16-P5-01-A | 14288 | Rep_3 | R3-T29 | - | - |
| JADAJL010000301.1 | SAMN16304027 | Res13-Abat-EA3-S5-02-A | pRes13-Abat-EA3-S5-02-A | 14288 | Rep_3 | R3-T29 | - | - |
| AFCZ02000003.1 | SAMN00114924 | OIFC032 | pOIFC032-101 | 101298 | Rep_3 | R3-T3 | - | EJP40261.1 |
| AFDM01000010.1 | SAMN00114925 | OIFC189 | pOIFC189-111 | 110967 | Rep_3 | R3-T3 | - | EJG16262.1 |
| AP022239.1 | SAMD00194616 | WP8-W18-ESBL-11 | pWP8-W18-ESBL-11_1 | 114365 | Rep_3 | R3-T3 | - | - |
| AYFW01000101.1 | SAMN02597401 | UH2107 | pABUH4-111 | 111007 | Rep_3 | R3-T3 | - | ETS68256.1 |
| CM008330.1 | SAMN04272870 | AC002-1-R4 | pAC0021R4 | 111165 | Rep_3 | R3-T3 | - | PCO03588.1 |
| CM009049.2 | SAMN08093361 | ZQ2 | p1ZQ2 | 110967 | Rep_3 | R3-T3 | - | PQJ03931.1 |
| CP004359.1 | SAMN02603104 | MDR-TJ | pABTJ2 | 110967 | Rep_3 | R3-T3 | - | AGG91013.1 |
| CP006769.1 | SAMN02641530 | ZW85-1 | ZW85p2 | 113866 | Rep_3 | R3-T3 | - | AHB93304.1 |
| CP010398.1 | SAMN03263969 | 6200 | p6200-114.848kb | 114848 | Rep_3 | R3-T3 | - | AJB68963.1 |
| CP010780.1 | SAMN03290686 | XH386 | pAB386 | 112157 | Rep_3 | R3-T3 | - | AKJ47876.1 |
| CP015484.1 | SAMN03277095 | ORAB01 | pORAB01-1 | 110965 | Rep_3 | R3-T3 | - | ANB90502.1 |
| CP016296.1 | SAMN04096368 | CMC-CR-MDR-Ab4 | pCMCVTAb1-Ab4 | 110968 | Rep_3 | R3-T3 | - | APQ87295.1 |
| CP016299.1 | SAMN04096369 | CMC-MDR-Ab59 | pCMCVTAb1-Ab59 | 110967 | Rep_3 | R3-T3 | - | APQ91163.1 |
| CP016301.1 | SAMN04096370 | CMC-CR-MDR-Ab66 | pCMCVTAb1-Ab66 | 110967 | Rep_3 | R3-T3 | - | APQ94950.1 |
| CP018257.1 | SAMN06077192 | AF-673 | pAF-673 | 110964 | Rep_3 | R3-T3 | - | APJ25221.1 |
| CP018333.1 | SAMN06099024 | A1296 | pA1296_1 | 112773 | Rep_3 | R3-T3 | - | ATI40464.1 |
| CP020585.1 | SAMN06650245 | CBA7 | pCBA7_1 | 111999 | Rep_3 | R3-T3 | - | ARG11462.1 |
| CP021327.1 | SAMN07135565 | XH386 | pXH386 | 112155 | Rep_3 | R3-T3 | - | AWW83408.1 |
| CP023025.1 | SAMN07520237 | 10324 | pAba10324c | 113420 | Rep_3 | R3-T3 | - | AXX47015.1 |
| CP024125.1 | SAMN07445112 | AYP-A2 | pAYP-A2 | 110967 | Rep_3 | R3-T3 | - | ATU25204.1 |
| CP026128.1 | SAMN06040401 | ABNIH28 | pABA-1fe1 | 110754 | Rep_3 | R3-T3 | - | AUT40134.1 |
| CP026945.1 | SAMN07977762 | S1 | pAbS1_02 | 111068 | Rep_3 | R3-T3 | - | AVG28499.1 |
| CP027182.1 | SAMN04014911 | AR_0070 | p4AR_0070 | 105127 | Rep_3 | R3-T3 | - | AVI35343.1 |
| CP027185.1 | SAMN04014893 | AR_0052 | p4AR_0052 | 105128 | Rep_3 | R3-T3 | - | AVI39456.1 |
| CP027483.1 | SAMN07689236 | I43 | pABI43 | 114918 | Rep_3 | R3-T3 | - | AVN23945.1 |
| CP028139.1 | SAMN08513260 | NCIMB 8209 | pAbNCIMB8209_134 | 133709 | Rep_3 | R3-T3 | - | QBC49395.1 |
| CP032217.1 | SAMN09906510 | UPAB1 | p2UPAB1 | 80061 | Rep_3 | R3-T3 | - | - |
| CP034093.1 | SAMN10441121 | A52 | pA52-1 | 110713 | Rep_3 | R3-T3 | - | QAB42408.1 |
| CP036284.1 | SAMN10261590 | TG60155 | p60155_1 | 127784 | Rep_3 | R3-T3 | - | QBH55712.1 |
| CP038645.1 | SAMN11311113 | ACN21 | p8ACN21 | 116047 | Rep_3 | R3-T3 | - | QBY91555.1 |
| CP039519.1 | SAMN10261589 | TG22653 | pTG22653 | 127784 | Rep_3 | R3-T3 | - | QCH35063.1 |
| CP041590.1 | SAMN12158047 | J9 | pJ9-3 | 145071 | Rep_3 | R3-T3 | - | QDM68418.1 |
| CP046537.1 | SAMN13476302 | XL380 | pXL380 | 112007 | Rep_3 | R3-T3 | - | QGW12467.1 |
| CP047974.1 | SAMN13884837 | DETAB-P2 | pDETAB1 | 103751 | Rep_3 | R3-T3 | - | QMS84036.1 |
| CP050905.1 | SAMN14308892 | DT-Ab057 | p2DT-Ab057 | 110967 | Rep_3 | R3-T3 | - | QIX32630.1 |
| CP050908.1 | SAMN14308866 | DT-Ab022 | p3DT-Ab022 | 119067 | Rep_3 | R3-T3 | - | QIX36501.1 |
| CP059387.1 | SAMN15541804 | 36-1512 | p1.36-1512 | 113330 | Rep_3 | R3-T3 | - | QLY88253.1 |
| CP061515.1 | SAMN12391854 | CFSAN093710 | pCFSAN093710_1 | 110967 | Rep_3 | R3-T3 | - | QNV19832.1 |
| CP061518.1 | SAMN12413929 | CFSAN093709 | pCFSAN093709 | 110965 | Rep_3 | R3-T3 | - | QNV16036.1 |
| CP061520.1 | SAMN12391536 | CFSAN093708 | pCFSAN093708 | 110967 | Rep_3 | R3-T3 | - | QNV35117.1 |
| CP061522.1 | SAMN12391534 | CFSAN093707 | pCFSAN093707 | 110964 | Rep_3 | R3-T3 | - | QNV31204.1 |
| CP061524.1 | SAMN12391535 | CFSAN093706 | pCFSAN093706 | 110967 | Rep_3 | R3-T3 | - | QNV27490.1 |
| CP062920.1 | SAMN16304032 | Res13-Abat-PEA21-P4-01-A | p4Res13-Abat | 113139 | Rep_3 | R3-T3 | - | QPF15204.1 |
| CP072527.1 | SAMN18498586 | DETAB-E227 | pDETAB4 | 113682 | Rep_3 | R3-T3 | - | QTM22032.1 |
| MK386680.1 | SAMN14228692 | ABAY04001 | pABAY04001_1A | 110967 | Rep_3 | R3-T3 | - | QBN23070.1 |
| CP050420.1 | SAMN14420254 | PM193665 | pPM193665_5 | 2762 | Rep_3 | R3-T30 | - | QJG86394.1 |
| CP050430.1 | SAMN14420255 | PM194188 | pPM194122_5 | 2762 | Rep_3 | R3-T30 | - | QJG82492.1 |
| GU978999.1 | - | - | p537 | 1125 | Rep_3 | R3-T31 | Aci5 | ADM89094.1 |
| LN865144.1 | SAMEA3449716 | CIP70.10 | pCIP70.10 | 7742 | Rep_3 | R3-T31 | Aci5 | CRL96367.1 |
| LN997847.1 | SAMEA3715145 | R2091 | pR2091 | 7742 | Rep_3 | R3-T31 | Aci5 | CUW37058.1 |
| GU979000.1 | - | - | p11921 | 1103 | Rep_3 | R3-T32 | Aci8 | ADM89095.1 |
| KY984047.1 | SAMN07509424 | AB242 | pAb242_25 | 24808 | Rep_3 | R3-T32 | Aci8 | AUO31913.1 |
| MG100202.1 | SAMN14228542 | Ab825 | pAb825_36 | 35743 | Rep_3 | R3-T32 | Aci8 | AVR61183.1 |
| CP033751.1 | SAMN10163233 | FDAARGOS_540 | p1FDAARGOS_540 | 7181 | Rep_3 | R3-T33 | - | AYX85221.1 |
| LR026974.1 | SAMEA4646219 | RDK39_49 | pKCRI-49-1 | 11681 | Rep_3 | R3-T33 | - | - |
| CP018334.1 | SAMN06099024 | A1296 | pA1296_2 | 11586 | Rep_3 | R3-T34 | - | ATI40537.1 |
| CP059304.1 | SAMN15574350 | AC1633 | pAC1633-3 | 9950 | Rep_3 | R3-T34 | - | QOJ62416.1 |
| AFDB02000003.1 | SAMN00114928 | Naval-81 | pNaval81-26 | 26089 | Rep_3 | R3-T35 | - | EJP56831.1 |
| ALII01000019.1 | SAMN02436485 | IS-123 | pIS123-18 | 17984 | Rep_3 | R3-T35 | - | EJO37647.1 |
| KY984047.1 | SAMN07509424 | AB242 | pAb242_25 | 24808 | Rep_3 | R3-T36 | - | AUO31910.1 |
| MG100202.1 | SAMN14228542 | Ab825 | pAb825_36 | 35743 | Rep_3 | R3-T36 | - | AVR61186.1 |
| CP000522.1 | SAMN02604331 | ATCC 17978 | pAB1 | 13408 | Rep_3 | R3-T37 | A1S_3471 | ABO13850.1 |
| CP000522.1 | SAMN02604331 | ATCC 17978 | pAB1 | 13408 | Rep_3 | R3-T37 | A1S_3471 | ABO13860.1 |
| CU468231.1 | SAMEA3138277 | SDF | p1ABSDF | 6106 | Rep_3 | R3-T38 | p1ABSDF0001 | CAP02936.1 |
| CU468231.1 | SAMEA3138277 | SDF | p1ABSDF | 6106 | Rep_3 | R3-T38 | p1ABSDF0001 | CAP02936.1 |
| CU468232.1 | SAMEA3138277 | SDF | p2ABSDF | 25104 | Rep_3 | R3-T39 | p2ABSDF0025 | CAP02966.1 |
| CU468232.1 | SAMEA3138277 | SDF | p2ABSDF | 25014 | Rep_3 | R3-T39 | p2ABSDF0025 | CAP02966.1 |
| AY541809.1 | SAMN14224286 | 19606 | pMAC | 9540 | Rep_3 | R3-T4 | Aci9 | AAT09649.1 |
| CM009036.2 | SAMN08093366 | ZQ7 | p1ZQ7 | 11191 | Rep_3 | R3-T4 | Aci9 | PST49955.1 |
| CM009653.1 | SAMN08093365 | ZQ6 | p5ZQ6 | 16476 | Rep_3 | R3-T4 | - | PQL72257.1 |
| CM012225.1 | SAMN07815360 | PIMB13AB-41 | pAB13-41 | 11194 | Rep_3 | R3-T4 | Aci9 | - |
| CP015122.1 | SAMN04621185 | ab736 | pab736 | 9539 | Rep_3 | R3-T4 | Aci9 | ARN32819.1 |
| CP024577.1 | SAMN07945345 | AbPK1 | pAbPK1a | 15113 | Rep_3 | R3-T4 | - | ATR89551.1 |
| CP027180.1 | SAMN04014911 | AR_0070 | p3AR_0070 | 73018 | Rep_3 | R3-T4 | Aci9 | AVI35133.1 |
| CP027186.1 | SAMN04014893 | AR_0052 | p2AR_0052 | 38505 | Rep_3 | R3-T4 | Aci9 | AVI39478.1 |
| CP035673.1 | SAMN07977426 | VB23193 | pVB23193 | 16033 | Rep_3 | R3-T4 | Aci9 | QBB78281.1 |
| CP035933.1 | SAMN10170272 | VB31459 | p1VB31459 | 11195 | Rep_3 | R3-T4 | Aci9 | QBF38536.1 |
| CP040086.1 | SAMN11571816 | VB33071 | p1VB33071 | 11194 | Rep_3 | R3-T4 | Aci9 | QCP44014.1 |

| | | | | | | | | |
|---|---|---|---|---|---|---|---|---|
| CP040088.1 | SAMN11571817 | VB35575 | pVB35575 | 11194 | Rep_3 | R3-T4 | Aci9 | QCP47688.1 |
| CP045109.1 | SAMN12389466 | ATCC 19606 | p2ATCC19606 | 9540 | Rep_3 | R3-T4 | Aci9 | QFQ03455.1 |
| CP065433.1 | SAMN08687991 | ATCC 17961 | pAB17961-1 | 9395 | Rep_3 | R3-T4 | Aci9 | QPP16179.1 |
| CP065886.1 | SAMN13450447 | FDAARGOS_917 | p2FDAARGOS_917 | 9540 | Rep_3 | R3-T4 | Aci9 | QQA24590.1 |
| LT594096.1 | SAMEA2439285 | BAL062 | pBAL062 | 8015 | Rep_3 | R3-T4 | Aci9 | SBS23985.1 |
| CU468233.1 | SAMEA3138277 | SDF | p3ABSDF | 24922 | Rep_3 | R3-T40 | p3ABSDF0002 | CAP02976.1 |
| CU468233.1 | SAMEA3138277 | SDF | p3ABSDF | 24922 | Rep_3 | R3-T40 | p3ABSDF0002 | CAP02976.1 |
| CU468233.1 | SAMEA3138277 | SDF | p3ABSDF | 24922 | Rep_3 | R3-T41 | p3ABSDF0009 | CAP02983.1 |
| CU468233.1 | SAMEA3138277 | SDF | p3ABSDF | 24922 | Rep_3 | R3-T41 | p3ABSDF0009 | CAP02983.1 |
| CU468233.1 | SAMEA3138277 | SDF | p3ABSDF | 24922 | Rep_3 | R3-T42 | p3ABSDF0018 | CAP02992.1 |
| CU468233.1 | SAMEA3138277 | SDF | p3ABSDF | 24922 | Rep_3 | R3-T42 | p3ABSDF0018 | CAP02992.1 |
| GU978996.1 | - | - | p736 | 1065 | Rep_3 | R3-T43 | Aci7 | ADM89091.1 |
| KT346360.1 | SAMN14226727 | RCH52 | pRCH52-1 | 11164 | Rep_3 | R3-T43 | - | ALC76579.1 |
| CP033753.1 | SAMN10163233 | FDAARGOS_540 | p3FDAARGOS_540 | 86551 | Rep_3 | R3-T44 | - | AYX85290.1 |
| CP044520.1 | SAMN12859885 | 29FS20 | p29FS20-1 | 66277 | Rep_3 | R3-T45 | - | QLF12435.1 |
| CP038259.1 | SAMN10386508 | 39741 | pEH_gr13 | 135229 | Rep_3 | R3-T46 | - | QBR82727.1 |
| CP042565.1 | SAMN12289292 | E47 | pE47_009 | 2427 | Rep_3 | R3-T47 | - | - |
| CP062923.1 | SAMN16304032 | Res13-Abat-PEA21-P4-01-A | p1Res13-Abat | 5242 | Rep_3 | R3-T48 | - | QPF15412.1 |
| AYFZ01000080.2 | SAMN02597404 | UH19608 | pABUH2a-5.6 | 5636 | Rep_3 | R3-T49 | - | ETQ55253.2 |
| GU978997.1 | - | - | p203 | 1068 | Rep_3 | R3-T5 | Aci3 | ADM89092.1 |
| CP021348.1 | SAMN03771402 | B8300 | pB8300 | 25150 | Rep_3 | R3-T5 | - | KMV24627.1 |
| CP038260.1 | SAMN10386508 | 39741 | pEH_gr3 | 25856 | Rep_3 | R3-T5 | - | QBR82828.1 |
| LR026972.1 | SAMEA4646212 | RDK36_28 | pKCRI-28-1 | 29606 | Rep_3 | R3-T5 | Aci3 | - |
| AFDL01000007.1 | SAMN04014897 | AR_0056 | pOIFC143-6.2 | 6241 | Rep_3 | R3-T5 | Aci3 | EJG16427.1 |
| CP003968.1 | SAMN02603576 | D1279779 | pD1279779 | 7416 | Rep_3 | R3-T5 | Aci3 | AGH37280.1 |
| CP007713.1 | SAMN02709859 | LAC-4 | pABLAC1 | 8006 | Rep_3 | R3-T5 | Aci3 | AIY39145.1 |
| CP018255.1 | SAMN06077191 | AF-401 | pAF-401 | 17583 | Rep_3 | R3-T5 | Aci3 | APJ21535.1 |
| CP018678.1 | SAMN05362953 | LAC4 | pALAC4-1 | 8006 | Rep_3 | R3-T5 | Aci3 | APO60695.1 |
| CP023024.1 | SAMN07520237 | 10324 | pAba10324b | 7143 | Rep_3 | R3-T5 | Aci3 | AXX46908.1 |
| CP040083.1 | SAMN11571814 | SP304 | pSP304 | 9185 | Rep_3 | R3-T5 | Aci3 | QCP40396.1 |
| CP050435.1 | SAMN14422682 | PM194229 | pPM194229_3 | 9847 | Rep_3 | R3-T5 | - | QJG78569.1 |
| CP065434.1 | SAMN08687991 | ATCC 17961 | pAB17961-2 | 6667 | Rep_3 | R3-T5 | - | QPP16194.1 |
| CP072529.1 | SAMN18498586 | DETAB-E227 | pDETAB6 | 7145 | Rep_3 | R3-T5 | Aci3 | QTM22127.1 |
| CP046900.1 | SAMN13565236 | A1429 | pA1429b | 19147 | Rep_3 | R3-T50 | - | QLB37617.1 |
| CM009050.2 | SAMN08093361 | ZQ2 | p4ZQ2 | 5695 | Rep_3 | R3-T51 | - | PQJ03811.1 |
| CP053216.1 | SAMN14833494 | DT0544C | p2DT0544C | 55394 | Rep_3 | R3-T52 | - | QLI38221.1 |
| CP030107.1 | SAMN09460321 | DA33382 | pDA33382-2-2 | 2372 | Rep_3 | R3-T53 | - | AXB17573.1 |
| CP040262.1 | SAMN11621520 | P7774 | p1P7774 | 5464 | Rep_3 | R3-T54 | - | QCR91187.1 |
| CP042564.1 | SAMN12289292 | E47 | pE47_008 | 3065 | Rep_3 | R3-T55 | - | QFH47722.1 |
| CP050408.1 | SAMN14414761 | VB2486 | pVB2486_5 | 5432 | Rep_3 | R3-T56 | - | QJH05174.1 |
| CP023023.1 | SAMN07520237 | 10324 | pAba10324a | 5300 | Rep_3 | R3-T57 | - | AXX46897.1 |
| CP034096.1 | SAMN10441121 | A52 | pA52-4 | 3610 | Rep_3 | R3-T58 | - | QAB42527.1 |
| AP023080.1 | SAMD00059694 | OCU_Ac16a | pOCU_Ac16a_3 | 13096 | Rep_3 | R3-T59 | - | - |
| CM009030.2 | SAMN08093369 | ZQ10 | p1ZQ10 | 35194 | Rep_3 | R3-T6 | - | PQJ03509.1 |
| CM009083.3 | SAMN08093368 | ZQ9 | p1ZQ9 | 35194 | Rep_3 | R3-T6 | - | PQJ03469.1 |
| CP021348.1 | SAMN03771402 | B8300 | pB8300 | 25150 | Rep_3 | R3-T6 | - | KMV24615.1 |
| CP038260.1 | SAMN10386508 | 39741 | pEH_gr3 | 25856 | Rep_3 | R3-T6 | - | QBR82859.1 |
| CP059478.1 | SAMN15637465 | 17-84 | p17-84_OXA | 108715 | Rep_3 | R3-T6 | - | QNB01606.1 |
| AYFH01000048.1 | SAMN02597386 | UH7607 | pABUH3a-8.2 | 8190 | Rep_3 | R3-T6 | - | ETR11568.1 |
| AYFZ01000083.1 | SAMN02597404 | UH19608 | pABUH3b-7.8 | 7819 | Rep_3 | R3-T6 | - | ETQ55058.1 |
| CM009035.2 | SAMN08093367 | ZQ8 | p3ZQ8 | 11034 | Rep_3 | R3-T6 | - | PQL85655.1 |
| CM009046.2 | SAMN08093363 | ZQ4 | p2ZQ4 | 13311 | Rep_3 | R3-T6 | - | PST50007.1 |
| CP033752.1 | SAMN10163233 | FDAARGOS_540 | p2FDAARGOS_540 | 13195 | Rep_3 | R3-T6 | - | AYX85239.1 |
| CP038648.1 | SAMN11311113 | ACN21 | p5ACN21 | 9205 | Rep_3 | R3-T6 | - | QBY91635.1 |
| CP042207.1 | SAMN08637743 | DS002 | pTS9900 | 9900 | Rep_3 | R3-T6 | - | QDX16364.1 |
| CP051868.1 | SAMN14667517 | Ab-C63 | pAb-C63_2 | 10663 | Rep_3 | R3-T6 | - | QJF41245.1 |
| CM009647.1 | SAMN08093361 | ZQ2 | p2ZQ2 | 12769 | Rep_3 | R3-T60 | | PQJ03830.1 |
| CP059302.1 | SAMN15574350 | AC1633 | pAC1633-4 | 5210 | Rep_3 | R3-T61 | - | QOJ62393.1 |
| KY617771.1 | SAMN14227876 | SGH0823 | pS30-1 | 18234 | Rep_3 | R3-T62 | - | ARH59503.1 |
| CP012956.1 | SAMN04029125 | D36 | pD36-4 | 47457 | Rep_3 | R3-T63 | - | ALJ89842.1 |
| CP053221.1 | SAMN14833556 | DT01139C | p1DT01139C | 9613 | Rep_3 | R3-T64 | - | QLI41769.1 |
| LR026973.1 | SAMEA4646218 | RDK37_43 | pKCRI-43-1 | 34935 | Rep_3 | R3-T65 | - | - |
| CP042559.1 | SAMN12289292 | E47 | pE47_003 | 8795 | Rep_3 | R3-T66 | - | QFH47692.1 |
| CP059303.1 | SAMN15574350 | AC1633 | pAC1633-2 | 12651 | Rep_3 | R3-T67 | - | QOJ62400.1 |
| KY984046.1 | SAMN07509424 | AB242 | pAb242_12 | 11891 | Rep_3 | R3-T68 | - | AUO31881.1 |
| GQ861437.1 | - | - | 135040 | 3975 | Rep_3 | R3-T69 | rep135040 | ACX70400.1 |
| CU459140.1 | SAMEA3138279 | AYE | p3ABAYE | 94413 | Rep_3 | R3-T7 | p3ABAYE0002 | CAM84695.1 |
| CU459140.1 | SAMEA3138279 | AYE | p3ABAYE | 94413 | Rep_3 | R3-T7 | p3ABAYE0002 | CAM84695.1 |
| CP027245.2 | SAMN08364584 | WCHAB005078 | pOXA58_005078 | 70509 | Rep_3 | R3-T7 | - | AVN12752.1 |
| CP033769.1 | SAMN10163228 | FDAARGOS_533 | p2FDAARGOS_533 | 97783 | Rep_3 | R3-T7 | p3ABAYE0002 | AYY55194.1 |
| CP038263.1 | SAMN10386510 | LEV1449/17Ec | pEC_gr13 | 128013 | Rep_3 | R3-T7 | p3ABAYE0002 | QBR79086.1 |
| CP041149.1 | SAMN12057668 | CUVET-MIC596 | pCUVET596 | 82016 | Rep_3 | R3-T7 | - | QJP32841.1 |
| CP042210.1 | SAMN08637743 | DS002 | pTS134338 | 134338 | Rep_3 | R3-T7 | p3ABAYE0002 | - |
| CP042558.1 | SAMN12289292 | E47 | pE47_002 | 59744 | Rep_3 | R3-T7 | p3ABAYE0002 | QFH47659.1 |
| CP044357.1 | SAMN12825295 | CAM180-1 | pCAM180A | 92034 | Rep_3 | R3-T7 | p3ABAYE0002 | QEY06053.1 |
| CP072528.1 | SAMN18498586 | DETAB-E227 | pDETAB5 | 97035 | Rep_3 | R3-T7 | - | QTM22087.1 |
| KT852971.1 | SAMN14226423 | 255_n | p255n_1 | 92939 | Rep_3 | R3-T7 | - | AMD83595.1 |
| AY228470.1 | - | - | pAB02 | 4162 | Rep_3 | R3-T8 | repA_ABE | AAR00517.1 |
| EU294228.1 | SAMN14225727 | nk | pABIR | 29823 | Rep_3 | R3-T8 | RepA_AB | ACB05788.1 |
| MN495625.1 | - | A2485 | pA2485 | 15405 | Rep_3 | R3-T8 | repA_ABE | QID24195.1 |
| MN495626.1 | - | A2503 | pA2503 | 15405 | Rep_3 | R3-T8 | repA_ABE | QID24215.1 |
| CP023021.1 | SAMN07520236 | 9201 | pAba9201a | 9024 | Rep_3 | R3-T8 | repA_ABE | AXX43430.1 |
| CP023027.1 | SAMN07520233 | 10042 | pAba10042a | 10062 | Rep_3 | R3-T8 | repA_ABE | AXX50757.1 |
| CP023035.1 | SAMN07520231 | 5845 | pAba5845a | 9935 | Rep_3 | R3-T8 | repA_ABE | AXX58329.1 |
| CP029572.1 | SAMN09241862 | DA33098 | pDA33098-9-2 | 8771 | Rep_3 | R3-T8 | repA_ABE | AWO18623.1 |
| GQ476987.1 | SAMN14225384 | CU2 | pMMCU2 | 10270 | Rep_3 | R3-T8 | repA_ABE | ACY68271.1 |
| GQ904226.1 | SAMN14225471 | - | pMMD | 9964 | Rep_3 | R3-T8 | repA_ABE | ADB23463.1 |
| KT022421.1 | SAMN14226656 | ML | pAB-ML | 12056 | Rep_3 | R3-T8 | repA_ABE | - |
| AYOI01000002.1 | SAMN02597423 | UH10707 | pABUH5-114 | 114115 | Rep_3 | R3-T9 | - | ETR59191.1 |
| CP026705.1 | SAMN04014897 | AR_0056 | tig00000058_pilon | 113706 | Rep_3 | R3-T9 | - | AVE44280.1 |
| CP026944.1 | SAMN07977762 | S1 | pAbS1_01 | 108394 | Rep_3 | R3-T9 | - | AVG28377.1 |
| CP027122.1 | SAMN04014897 | AR_0056 | p3AR_0056 | 113706 | Rep_3 | R3-T9 | - | AVN03837.1 |
| CP027608.1 | SAMN04014943 | AR_0102 | p2AR_0102 | 106967 | Rep_3 | R3-T9 | - | - |
| CP029570.1 | SAMN09241862 | DA33098 | pDA33098-108 | 108151 | Rep_3 | R3-T9 | - | AWO18398.1 |
| CP031445.1 | SAMN09769497 | MDR-UNC | pMDR-UNC | 112216 | Rep_3 | R3-T9 | - | QBA07831.1 |
| CP035046.1 | SAMN05238672 | ABUH793 | p107.0Kbp | 106963 | Rep_3 | R3-T9 | - | QAS99596.1 |
| CP040426.1 | SAMN11660471 | PB364 | pPB364_1 | 111449 | Rep_3 | R3-T9 | - | QCT18072.1 |
